# High-throughput single-molecule quantification of individual base stacking energies in nucleic acids

**DOI:** 10.1101/2022.05.25.493108

**Authors:** Jibin Abraham Punnoose, Kevin J. Thomas, Arun Richard Chandrasekaran, Javier Vilcapoma, Andrew Hayden, Kacey Kilpatrick, Sweta Vangaveti, Alan Chen, Thomas Banco, Ken Halvorsen

## Abstract

Base stacking interactions between adjacent bases in DNA and RNA are known to be important for many biological processes, for drug development, and in other biotechnology applications. While previous work has estimated base stacking energies between pairs of bases, the individual contributions of each base to the stacking interaction has remained unknown. Here, we developed a novel methodology using a custom Centrifuge Force Microscope to perform high-throughput single molecule experiments to measure base stacking energies between individual adjacent bases. We found stacking energies strongest between purines (G|A at −2.3 ± 0.2 kcal/mol) and weakest between pyrimidines (C|T at −0.4 ± 0.1 kcal/mol). Hybrid stacking with phosphorylated, methylated, and RNA bases had no measurable effect, but a fluorophore modification reduced stacking energy. The implications of the work are demonstrated with three applications. We experimentally show that base stacking design can influence assembly and stability of a DNA nanostructure, modulate kinetics of enzymatic ligation, and determine accuracy of force fields in molecular dynamics (MD) simulations. Our results provide new insights into fundamental DNA interactions that are critical in biology and can inform rational design in diverse biotechnology applications.

## Main Text

DNA is remarkable in its ability to efficiently carry genetic information, and for material properties that provide high overall stability and still allow biological manipulation. These features are governed primarily by base pairing between two complementary strands and coaxial base stacking between adjacent bases. Both play important roles in nucleic acid structure and function, but base stacking interactions are sometimes overlooked. An interesting example is a minimal RNA kissing complex, with only 2 canonical base pairs but unusually high mechanical stability (similar to a ~10 bp duplex) [1] attributed largely to base stacking interactions [2,3]. Indeed, base stacking is critical to biological processes including DNA replication [4,5], RNA polymerization [6], and formation and management of G-quadruplexes in telomeres [7,8]. Base stacking is also thought to be critical for supramolecular assembly of nucleobases in pre-biotic RNA as part of the RNA world hypothesis [24,25]. Stacking also affects drug development, since small molecule intercalators targeting DNA or RNA rely on stacking interactions to disrupt a multitude of diverse diseases including cancers, viral infections, Myotonic dystrophy, and Parkinson’s disease [9–11]. In biotechnology, synthetic base analogs such as LNA [12], universal bases [13], and size expanded bases [14] partly rely on modified base stacking interactions. The formation of synthetic DNA nanostructures can rely heavily on base stacking, including DNA polyhedra [15], and DNA crystals [16] and liquid crystals [17], with some designs assembling using only blunt end stacking interactions [18,19].

Measuring stacking energy between adjacent bases in a helix is challenging due to the small energies, the difficulty in disentangling base pairing and base stacking contributions, and experimental limitations. Early studies used thermal melting spectrophotometry with different terminal overhanging ends to resolve these effects [26,27]. More recent direct experimental studies of stacking interactions used polyacrylamide gel electrophoresis (PAGE) assays of nicked dsDNA to quantify pairs of stacking interactions [20,21], or optical tweezers to monitor binding and unbinding of DNA nanobeams with terminal stacking interactions [22]. These studies have made immense contributions to our knowledge, but their designs and experimental constraints precluded the measurement of base stacking between two individual bases rather than pairs of bases.

Here we set out to directly measure individual base stacking interactions at the single molecule level. Single-molecule pulling techniques can apply biologically relevant picoNewton-level forces to individual molecules, and have been indispensable for the study of biomolecules including folding dynamics and mechanisms of biomolecular interaction [28]. Force is a useful perturbation to compare bond strengths, enabling faster dissociation while still allowing quantification of solution (force-free) behavior. Common single-molecule methods that apply force include optical and magnetic tweezers and atomic force microscopy (AFM). We expanded the single-molecule toolkit with the development of the Centrifuge Force Microscope (CFM), a high-throughput technique that combines centrifugation and microscopy to enable many single-molecule force-clamp experiments in parallel [23]. We have made several iterations to improve the technique, notably enabling single-molecule manipulation with a benchtop centrifuge [29–31], and other groups have advanced the technique as well [32,33]. The high throughput nature of the CFM makes it well suited to collect data from thousands of pulling experiments for a comprehensive assessment of individual base stacking interactions.

Combining the high-throughput CFM with novel DNA construct design we quantified individual stacking energies of 10 unique base combinations ranging from −2.3 ± 0.2 kcal/mol (G|A stack) to −0.4 ± 0.1 kcal/mol (C|T stack). Stacking energy was not measurably affected by phosphorylation, methylation, or substitution by an RNA nucleotide, but was reduced by a bulky fluorophore modification. Applying our results, we used base stacking to alter the structural stability of a DNA tetrahedron and to change the kinetics of an enzymatic ligation reaction. We also show that our results can be used to evaluate accura y and potentially improve force fields in MD simulations. Our work represents the first comprehensive picture of individual base stacking interactions, and provides concrete examples of how such knowledge can be applied.

## Results

### Premise of experimental design

Base stacking interactions (Fig. 1a) are relatively weak on the order of ~1 kcal/mol, making the measurement of individual stacking interactions challenging. To address this, we considered the design of two duplexes that are weakly held together by identical base pairing but differ by presence or absence of a terminal base stack (Fig. 1b). This terminal base stack strengthens the interaction and lowers the energy of the bound state (Fig. 1c). The application of external force shifts the process out of equilibrium, allowing only the bound to unbound transition. Measurement of dissociation kinetics can then be used to determine the effect of a single terminal base stack (Fig. 1d). This design allows for flexibility in the overall experimental time scale by control of both the design of the base pairs in the central duplex and by the magnitude of the externally applied force. Building from previous work where we resolved the energy difference of a single nucleotide polymorphism [31], we hypothesized that properly designed single-molecule pulling experiments could resolve the solution-based energies of individual base stacking interactions.

**Figure 1:**
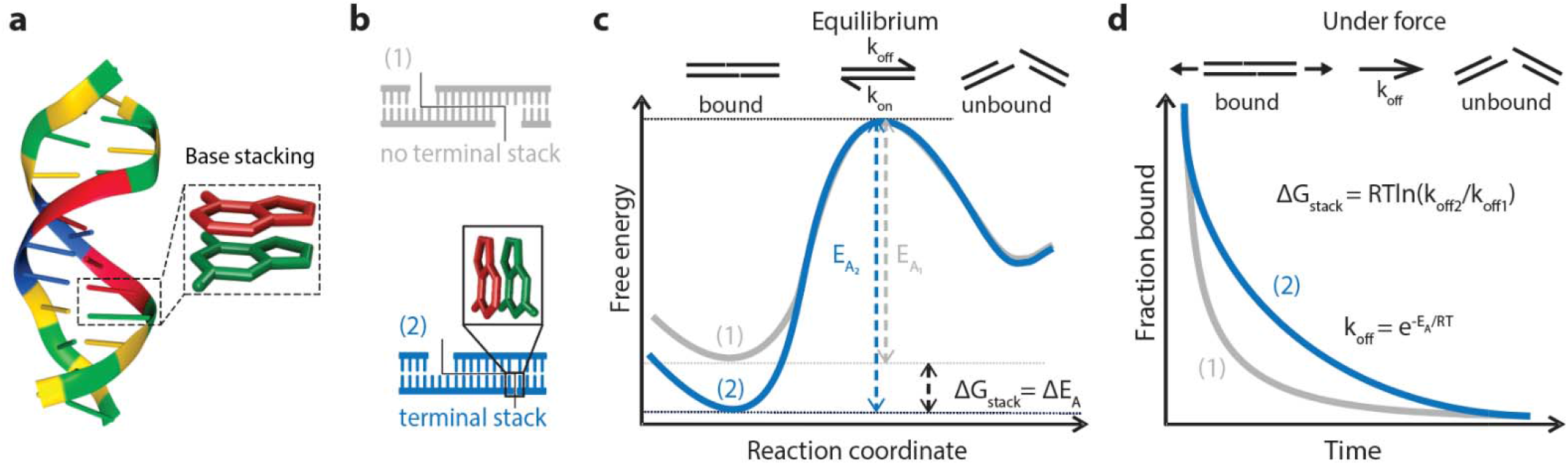
Conceptual overview. (a) Model of a DNA duplex [34] with enlarged frame showing stacked adjacent bases. (b) Design of two duplexes differing by a single base stacking interaction. (c) Free-energy diagram of a DNA duplex with and without a terminal base stack. The base stack primarily increases the activation energy from E_A1_ to E_A2_, with the difference representing the free energy of the single terminal base stack, ΔG_stack_. (d) External force lowers the activation barriers and prevents rebinding, causing exponential dissociation that can be experimentally measured and used to calculate stacking free energy.

To enable high throughput single-molecule pulling experiments, we used a custom designed CFM. The CFM is essentially a microscope that can be centrifuged, providing a controlled force application to single-molecule tethers, coupled with video microscopy imaging that can track individual tethers during the experiment (Fig. 2a). Using advances in 3D printing, cameras, and wireless communication electronics, we recently integrated the microscope into a bucket of a standard benchtop centrifuge (Fig. 2b). We achieved live streaming of microscopy images during centrifugation by WiFi communication with an external computer that controls both the centrifuge and the camera through custom Labview software (Fig. 2c). During a typical experiment, we observe tens to hundreds of tethered microspheres in a full field of view at 40x magnification (Fig. 2d). As the centrifuge spins, force is applied to a DNA construct tethered between a glass slide and microspheres, forcing dissociation of the duplex over time and causing the microspheres to disappear from view (Fig. 2e). Each microsphere is monitored to track individual dissociation events (Fig. 2f), which are used to create a dissociation curve that can be used to extract the off rate (Fig. 2g).

**Figure 2:**
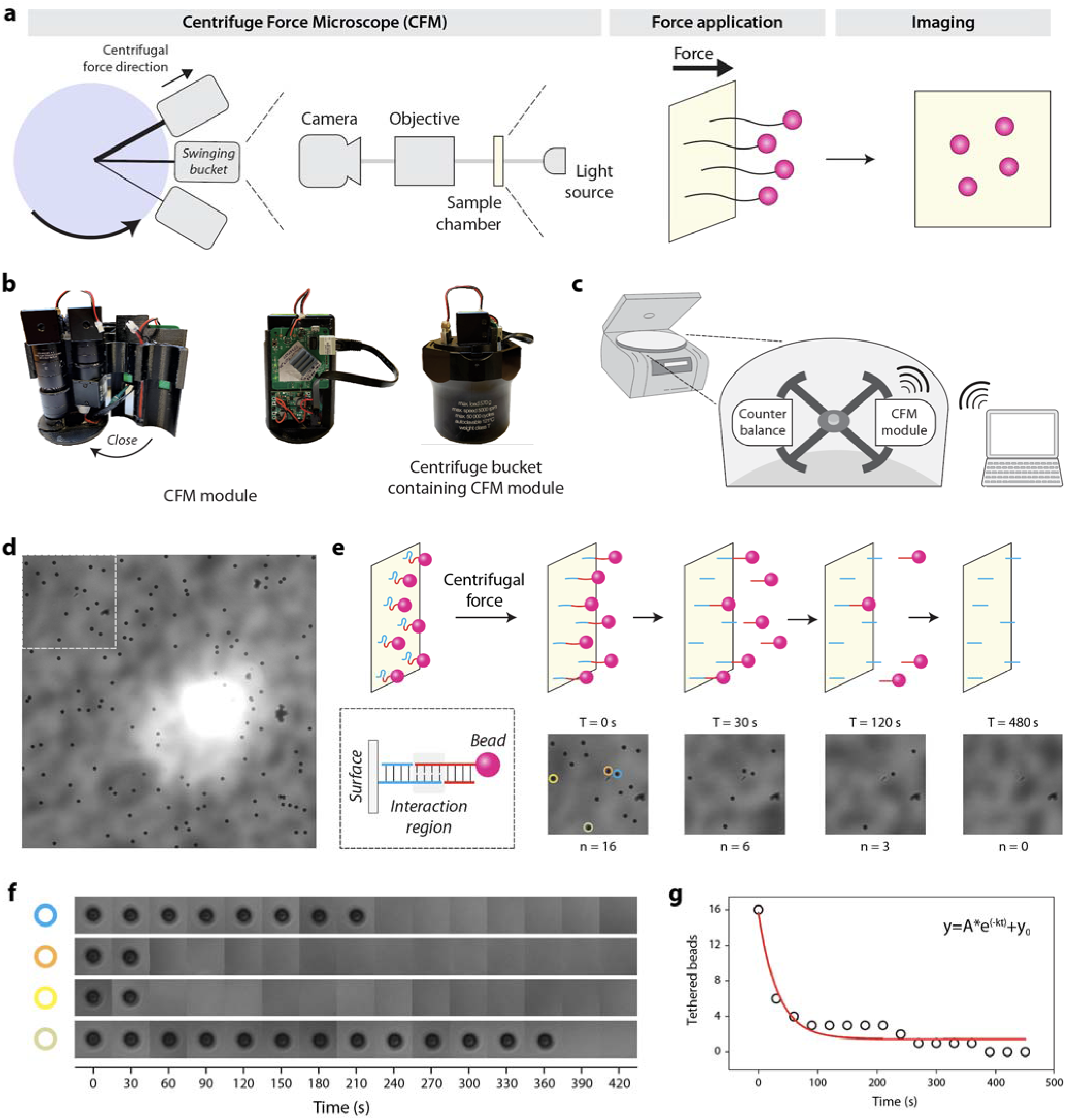
Concept of the Centrifuge Force Microscope (CFM) and force clamp assay. (a) The CFM is comprised of a video microscope that is centrifuged. Centrifugal force is applied to tethered microspheres and aligns with the imaging pathway to give a head on view of microspheres. (b) Images of the custom CFM module show the compact central optics, a clamshell style 3D printed housing, and supporting electronics, which fit inside a centrifuge bucket. (c) The CFM module operates in a benchtop centrifuge, which is controlled by an external computer that receives a live video stream by WiFi. (d) A typical microscopy image of ~100 tethered beads at a ~40x magnification. (e) Concept and partial-frame images of tether dissociation observed in the force clamp assay. As the weak central duplex dissociates, tethered beads fall out of focus and disappear from view. (f) Custom MATLAB software tracks tethered beads over time and records dissociation times. Four examples shown correspond to a subset of beads in panel (e). (g) Decay plot obtained from the dissociation time analysis of the tethers in sub-frame (e). The red line is a single-exponential fit to extract off rate.

### Experimental measurement of single base stacking energies

As a first test, we confirmed that kinetic differences were measurable between DNA constructs varying by a single base stacking interaction. We designed and created three DNA constructs with identical short central duplexes (8 bp) but varying terminal base stacking interactions (Fig. 3a and Fig. S1). In the control construct, a 3 nt poly-T spacer was used to eliminate terminal base stacking completely. Unlike most previous designs that look at pairs of base stacks or groups of pairs, this design isolates the contribution of a single base stacking interaction between two individual bases. We adopt a notation of X|Y to indicate a stacking interaction between bases X and Y read in the 5’ to 3’ direction, with X residing on the 3’ end of one strand and Y on the 5’ of another. Constructs were created by self-assembly of the 7249 nt M13 genomic ssDNA with complementary tiling oligonucleotides, similar to our previous work with DNA nanoswitches [35,36]. The oligonucleotides tile along the length to make double stranded DNA, to provide a terminal double biotin for coupling to surfaces, and to provide “programmable” overhanging ends comprising half of the central duplex (sequences in Table S1). For the experiment, two pairing DNA constructs were attached separately by biotin-streptavidin interactions to the microspheres and the cover glass. The microspheres were briefly allowed to come into contact with the cover glass within the reaction chamber to allow tethers to form before applying force by centrifugation and measuring dissociation.

**Figure 3:**
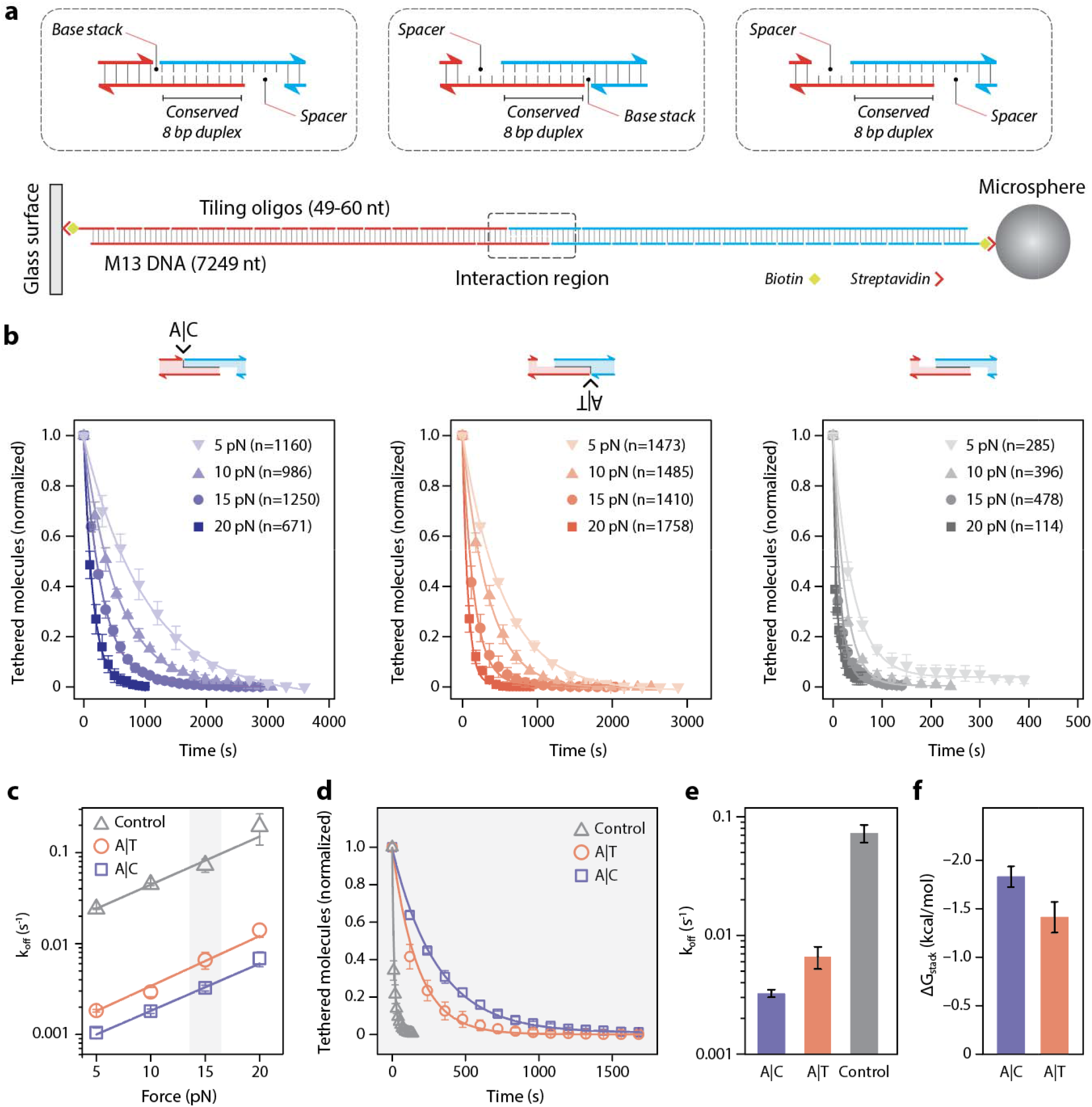
Experimental measurement of single base stacking energies. (a) A weak central 8 bp duplex is designed to be flanked by a terminal base stack or no base stacks. The central interaction is formed between two DNA handles attached to a glass slide and a microsphere through biotin-streptavidin interaction. (b) Raw data and single exponential fits obtained for the A|C, A|T and control constructs at forces from 5-20 pN. Error bars represent standard deviation from at least three replicates. (c) Force-dependent off-rates fit with the Bell-Evans model (solid lines) to determine thermal off-rate. Error bars represent standard deviation in off-rates from individual replicates (Figure S2-S4). (d) Analysis of the three constructs at 15 pN shows clear differences in dissociation, fit with exponential decay curves to yield off-rates (e), from which ΔG_stack_ is calculated (f).

We probed the duplexes at forces from 5-20 pN to establish force dependent dissociation rates at room temperature (21 ± 1 °C). We hypothesized that the characteristic force scale of different constructs should be nearly identical, which would allow us to extract equilibrium energy differences from off-rates obtained at any constant force. We collected data from over 10,000 single-molecule tethers from multiple experiments that ranged from a few minutes to an hour to ensure most or all beads were dissociated (Fig. 3b). The data were well described by single exponential decays to determine off-rates at different forces (Fig. S2-S4). Using the Bell-Evans model [37,38], we fit a linear trend to the logarithm of the force dependent off-rates for single A|C or A|T stacks and the no-stack control (Fig. 3c). We observed that force-dependent off rates were easily distinguishable between the constructs but followed identical slopes. This result confirmed that individual base stacking interactions could be measured with this approach, and that the choice of force should not appreciably affect the calculated values of equilibrium free-energy of stacking. Using the 15 pN force as an example, it is clear that the three measurements are distinctly different (Fig. 3d,e), enabling the calculation of ΔG_stack_ by the ratio of off rates (Fig. 3f). We also verified consistency in calculated ΔG_stack_ across force values and found all results overlapping within error estimates (Fig. S5). We decided to proceed with a force of 15 pN, enabling centrifuge runs with hundreds of individual single-molecule experiments to complete in the 10-100 minutes time scale.

Having successfully proven the concept, we aimed to measure base stacking interactions between all four canonical bases (A,G,C,T) in DNA. Neglecting directionality, the four bases give rise to ten unique base stacks T|T, C|T, A|T, G|T, C|C, A|C, G|C, A|A, G|A, and G|G. Following our validated approach, we designed DNA constructs to isolate the effect of a single base stack for all 10 combinations (Fig. S6). To accomplish this with minimal disturbance to the central duplex, we designed the central duplex to have A,C,T, and G as the 4 terminal bases. This design allowed manipulation of the strands to accomplish all of the 10 combinations with only two control constructs (oligonucleotides listed in Tables S1-S3).

For each construct and control, we ran experiments at 15 pN at room temperature until most beads have dissociated. Each condition was run with at least three experimental replicates, where each run also contained tens to hundreds of individual tethers. We collected and analyzed over 10,000 single molecule tethers to measure the 10 base stacking interactions. From the images of each run, we measured the dissociation time for each molecule, constructed the decay plot, and found the off-rate by fitting with a single-exponential decay (Fig. 4a-b, Fig. S7-S9). We determined base stacking energies for all ten base stacks, ranging from −2.3 ± 0.2 kcal/mol for G|A (the strongest) to −0.4 ± 0.1 kcal/mol for C|T (the weakest) (Fig. 4c,Table 1). We observed a general trend that stacking energetics follows the order purine-purine > purine-pyrimidine > pyrimidine-pyrimidine. It is interesting to note that the two control constructs had nearly identical off-rates even with a 5’ to 3’ reversal of the central duplex.

**Figure 4:**
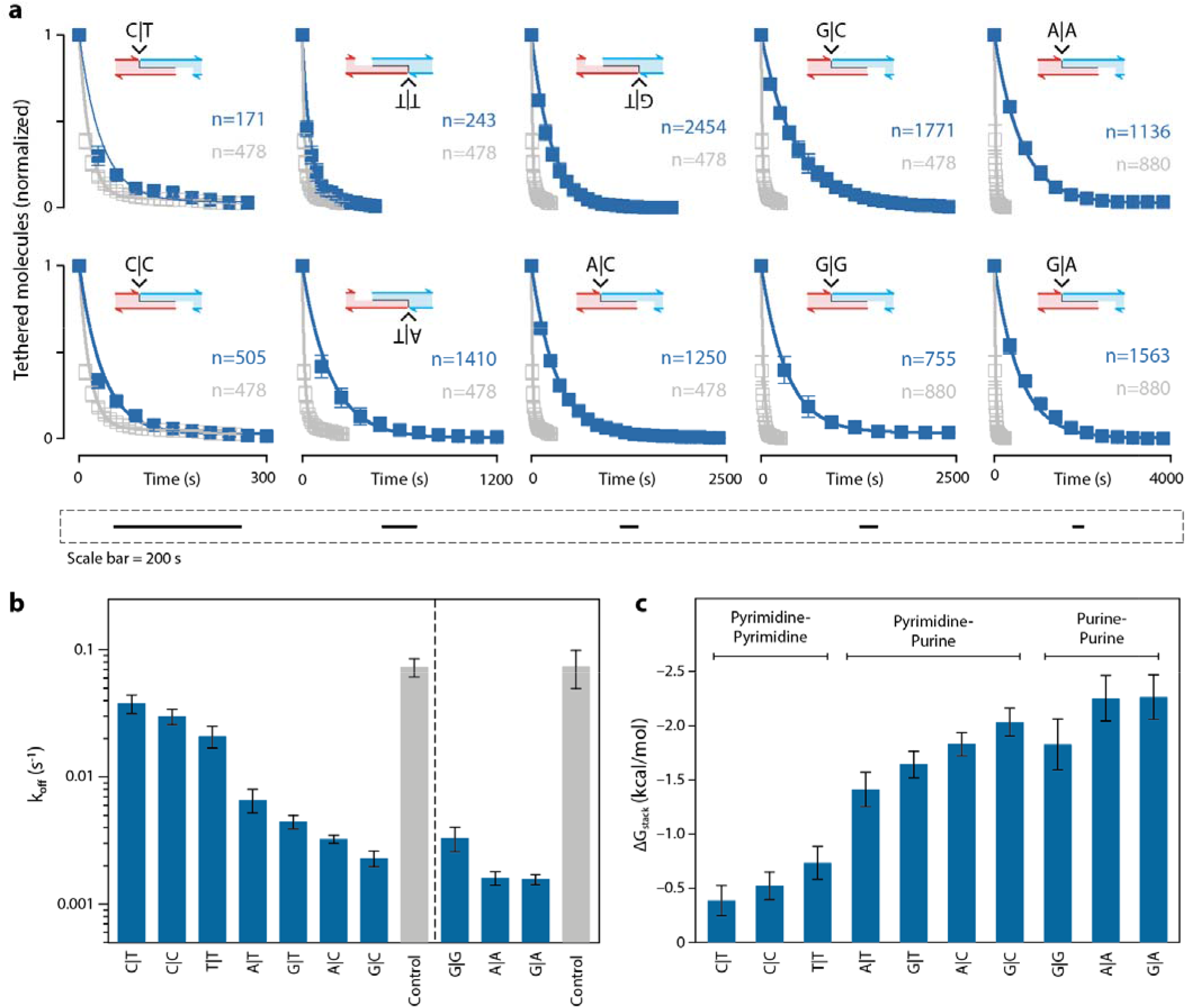
Comprehensive study of DNA base stacking. (a) Decay curves and single exponential fits obtained for unique base stacking combinations and their controls at a constant force of 15 pN. Error bars represent standard deviations from three data sets each consisting of one or more force clamp experiments. (b) Off-rates of DNA tethers containing various base stacks and their corresponding controls. Error bars represent standard deviation in off-rates from individual data sets (Fig. S7-S9). (c) Base stacking energies calculated from panel (b). Error bars represent calculated error propagation of results in (b).

**Table 1:** Individual base stacking energies determined using CFM.

|  | G\|A | A\|A | G\|G | G\|C | A\|C | G\|T | A\|T | T\|T | C\|C | C\|T |
| --- | --- | --- | --- | --- | --- | --- | --- | --- | --- | --- |
| $\Delta G_{stack}$<br>(kcal/mol) | $-2.3 \pm 0.2$ | $-2.3 \pm 0.2$ | $-1.8 \pm 0.2$ | $-2.0 \pm 0.1$ | $-1.8 \pm 0.1$ | $-1.6 \pm 0.1$ | $-1.4 \pm 0.2$ | $-0.7 \pm 0.2$ | $-0.5 \pm 0.1$ | $-0.4 \pm 0.1$ |

### Influence of nucleotide modification on base stacking energy

Various chemical modifications on nucleotides can influence base stacking and base pairing energy thereby affecting the stability of nucleic acid structures [39]. We extended our approach to probe the effect of these types of modifications in comparison with canonical bases. In particular, we chose phosphorylation, methylation, fluorescein (6-FAM) and substitution of deoxyribose to ribose to study their impact on stacking of the A|C base stack (Fig. 5a). We designed modified oligonucleotides and constructed four duplexes with modified A|C stacks and individual no-stacking controls for each modification (Fig. S10). Analogous to the regular base stacking experiments, we performed 15 pN force clamps and fit decay plots to obtain the off-rates (Fig. 5b-c, Fig S11-S12) used to calculate the stacking energy of the modified A|C base stack. The control constructs were all found to be consistent within error. We observed that phosphorylation, methylation and hybrid DNA-RNA stacks are not appreciably different from the regular A|C base stack, while the bulky FAM group reduced the base-stacking energy by 0.7 ± 0.1 kcal/mol (Fig. 5d). These results show that stacking effects of chemical modifications can be measured with our technique, and suggest generally that small modifications are less likely to interfere with stacking under the conditions tested here.

**Figure 5:**
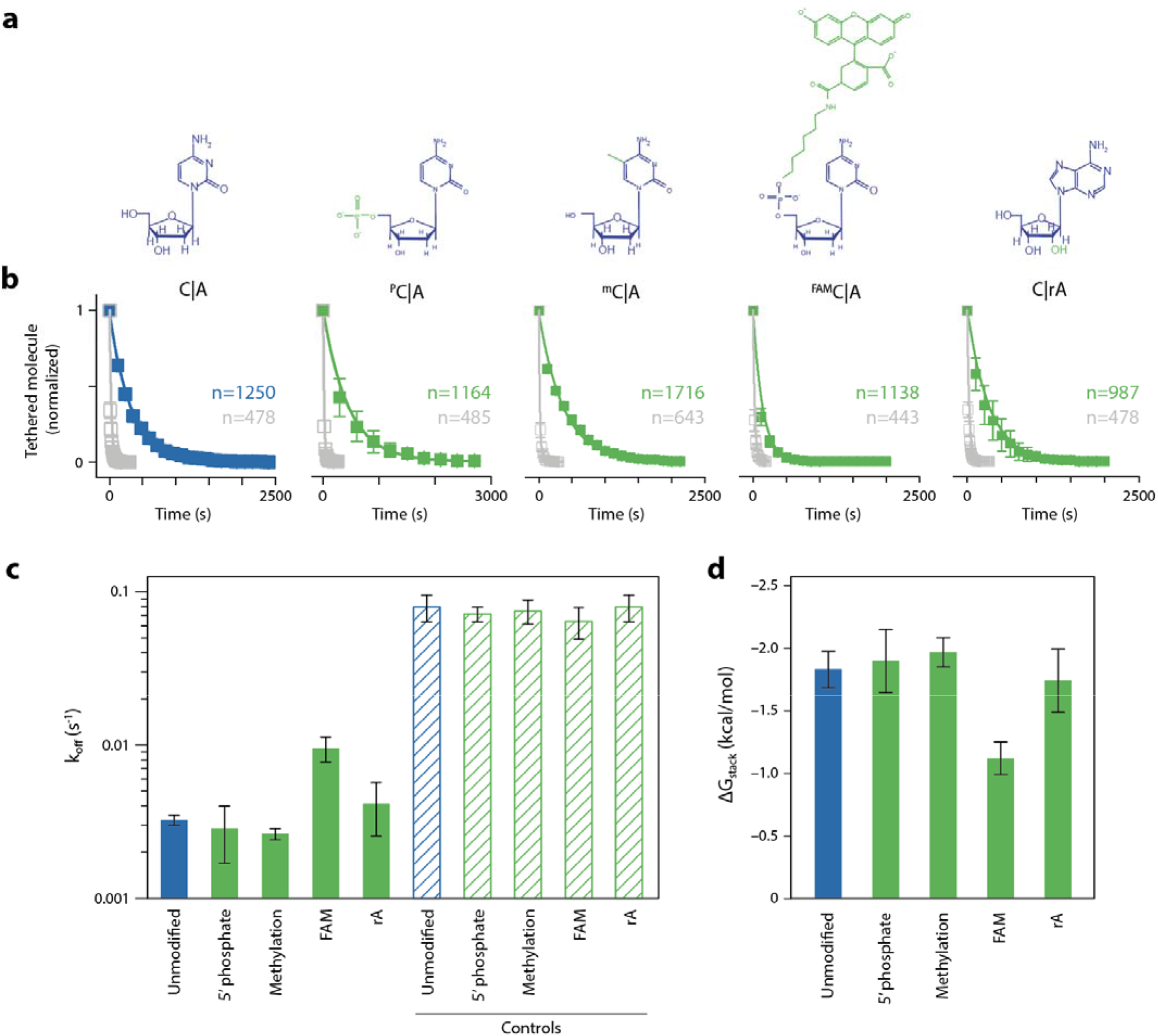
Effect of nucleotide modification on base stacking energy of nucleotides. (a) Modifications used in the study including 5′ Phosphorylated C, 5-methyl C, 5′ FAM modified C, 3′ ribose A. (b) Experimental data shows dissociation over time for constructs relative to their controls. Error bars represent standard deviation of triplicate data sets (c) Off-rates observed for tethers with modified bases and their controls. Error bars represent standard deviation in off-rates from individual data sets (Fig. S11-S12). (d) Free-energy of stacking calculated from off-rates in (c), with error bars propagated from results in (c).

### Base-stacking in biotechnology applications

Elucidation of these base stacking energies can benefit many aspects of biotechnology, which often rely on forming or dynamically controlling short DNA duplexes. These include molecular biology methods, such as genetic recombination, polymerase chain reaction, and sequencing, as well as emerging technologies like gene editing, synthetic biology, and DNA nanotechnology. These stacking energies can also help influence molecular simulations, whose accuracy relies on parameters that reflect realistic potentials between the simulated components. Here we show how our results can be used to benefit DNA nanotechnology, enzymatic ligation, and molecular dynamics (MD) simulations.

In DNA nanotechnology, DNA is used as a building block for nanomaterials [40] with applications including drug delivery [41] and sensing [42]. The field relies on forming controlled contacts between short DNA segments. We hypothesized that designing DNA motifs with specific interfacial base stacks could alter the assembly and stability of DNA nanostructures. To test this, we used the DNA tetrahedron as a model system, a widely used structure with biosensing and drug delivery applications [15,43]. The DNA tetrahedron is hierarchically self-assembled from 3-point-star motifs that connect to each other through a pair of 4 nt sticky ends (Fig. 6a). Our control structure, based on a DNA tetrahedron we previously reported [44], contained two pairs of base stacks (G|A and A|T) across two 4-nt sticky end connections. We annealed the DNA tetrahedron and validated self-assembly using non-denaturing polyacrylamide gel electrophoresis (PAGE) (Fig 6b). To test the effect of different base stacking interactions, we modified the sequence of the component DNA strands and constructed three other versions of the DNA tetrahedron with one pair of G|A base stacks, one pair of A|T base stacks, or no base-stacks (Fig. 6c). We observed that structures containing both the G|A and A|T bases stacks were best formed, followed by the G|A structure, while the other two were apparently too weak to form stable structures (Fig. 6d, full gels in Fig. S14). To confirm that the G|A design was less stable and not just produced in a lower quantity, we tested thermal stability and observed a decrease in the relative stability of the structures with increased temperature, and a clear indication that the G|A structure was unstable at 40° C while the G|A + A|T structure was still intact (Fig. 6e, Fig. S14). These results are consistent with our findings that G|A base stack is stronger than A|T base stack, demonstrate the crucial role of base-stacking in the stability of DNA nanostructures, and show for the first time how altered stability of a DNA tetrahedron can be achieved by design of base stacking interactions. Designing DNA nanostructures typically only involves consideration of the base pairing, and this work points to a new dimension of control and design flexibility.

**Figure 6:**
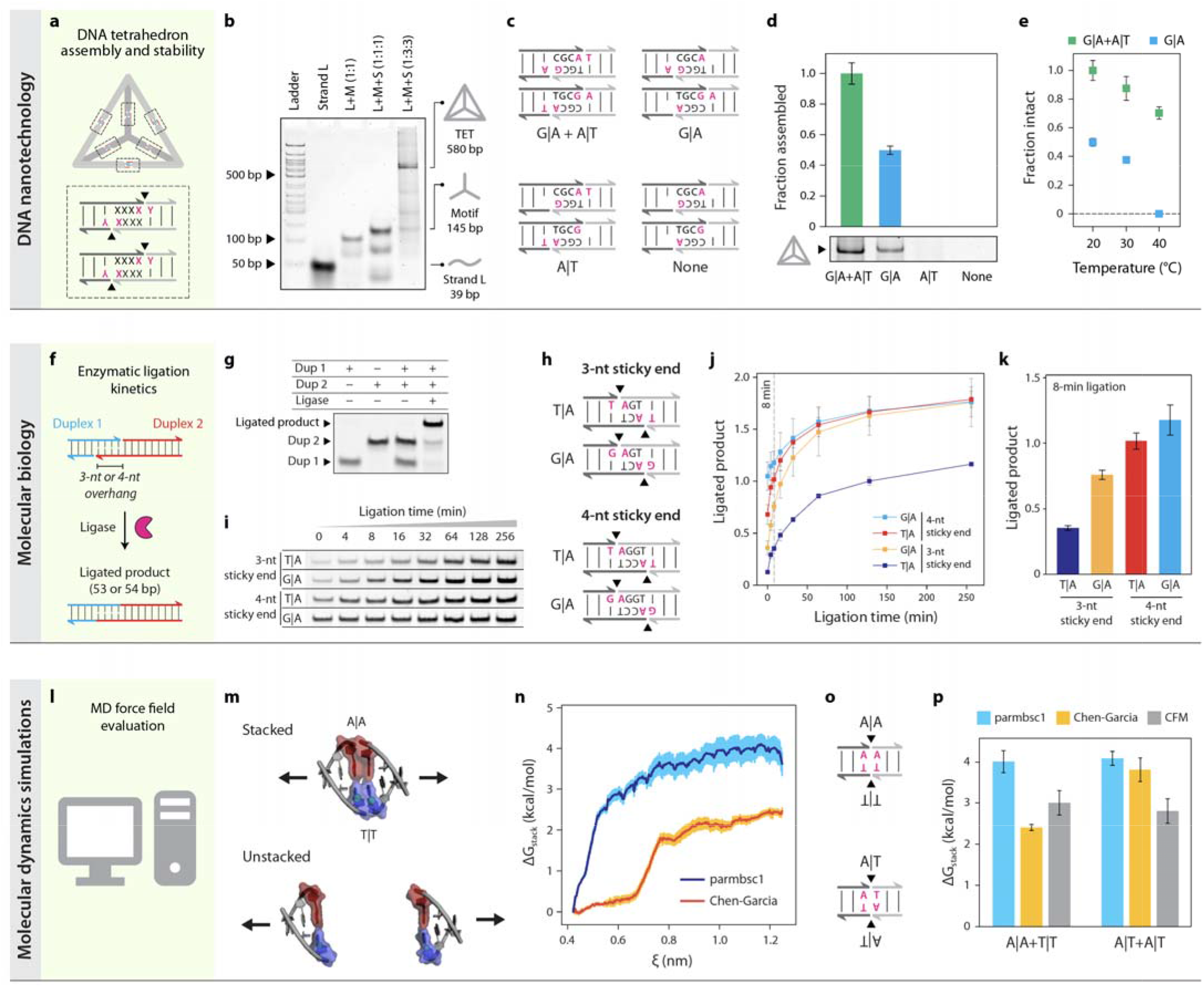
Base-stacking in biotechnology applications. (a) A DNA tetrahedron is assembled from four 3-point-star DNA motifs connected on each edge with two pairs of sticky ends. (b) Non-denaturing PAGE analysis shows DNA tetrahedron assembly from oligonucleotide components. (c) Designs of base stacking interactions tested in DNA tetrahedra that conserve sticky end sequences. (d) Assembly of DNA tetrahedra with different base stacks. (e) Thermal stability of DNA tetrahedra with various base-stacks. (f) Ligation of two DNA duplexes with 3 or 4 base pair sticky ends. (g) Non-denaturing PAGE confirms the ligation of the two DNA duplexes. (h) Designs of base stacking interactions of sticky ends tested for ligation. (i) Gel images show the increase in band intensity of ligated fragments (full gels in Fig. S15-16). (j) Quantified ligation product over time. (k) Ligated product for different base stacks at 8 minutes. (l) Molecular dynamics simulations of base stacking interactions with different force fields, (m) Simulation scheme showing two 3-mer duplexes with A|A in red and T|T in blue, in the initial (stacked) and the final (unstacked) confirmation. The pulling force is applied on the C1’ atoms of the T|T stacked pair, orthogonal to the base-pairs (n) Potential of mean force (PMF) of the A|A-T|T construct as a function of the distance between the pull groups (ξ), for the Amber99-Chen-Garcia [50] and parmbsc1 [52] force-fields. (o) Designs of base stacking interactions tested in MD simulation (p) Free energy of stacking (ΔG_stack_) as calculated from the simulations compared to experimentally determined values (values added from Table 1). Error bars in (d,e,j,k) represent standard deviation from triplicate experiments, in (n) represent the standard deviation for potentials sampled at each distance, and in (p) represent the average of standard deviations for potentials sampled at distance zeta > 1.1 nm.

DNA ligation is a process of enzymatically joining two pieces of DNA, often facilitated by short “sticky ends” of 1-4 nt that hybridize together. Ligation is a fundamental biological process that is required for DNA repair and replication and is integral to a wide range of biotechnology applications including sequencing, cloning, and diagnostics [44,45]. We hypothesized that modification of interfacial base stacks could alter the kinetics by changing the lifetime of the bound duplex and potentially the final efficiency of enzymatic ligation. To test this hypothesis, we designed and created short duplexes to enable ligation of products with varying sticky ends (Fig. 6f). First, we validated construction of the individual duplexes and the successful ligation of the two duplexes (Fig 6g). Next we investigated the ligation kinetics of four variants, 4 nt and 3 nt sticky ends with either T|A or G|A terminal base stacks (Fig. 6h,i and Fig. S15-S16). In the 4 nt case, we observed a slight increase in kinetics with the G|A stacks, which was most evident in the first 20 minutes of the reaction (Fig. 6j). For the 3 nt case, the difference was more striking, with a substantial difference in both the kinetics of ligation as well as the endpoint. The differences can be most clearly seen when looking at the ligated products after an 8 minute reaction, where the trend follows 4 nt G|A > 4 nt T|A > 3 nt G|A > 3 nt T|A (Fig. 6k). Interestingly, the magnitude of the change between T|A and G|A in the 3 nt case is similar to the change between 3 nt and 4 nt in G|A, suggesting that strong base stacking interactions could potentially compensate for weak base pairing in such short duplexes. Building on this idea, we tested whether we could design a 3 bp interaction with strong base stacking that outperforms a 4 bp interaction with weak base stacking. We made a 3 bp design with A|G stacks on both sides and found that it had substantially faster ligation kinetics than the same sequence with an added A-T pair but with two weaker C|T stacks (Figure S17). These results clearly show how our data can be used in biotechnology applications, presumably for a host of enzymatic interactions that go far beyond ligation.

Molecular dynamics (MD) simulations are a powerful tool to study conformational dynamics of biomolecules including nucleic acids [46]. However, simulations are only as accurate as their underlying empirical energy functions (i.e. force-fields), which must be strategically calibrated against experimental measurements. Force fields for nucleic acids require precise and separate calibration of base-stacking and base-pairing energies for all nucleotide combinations, which is particularly difficult to compare with experimental studies that typically combine these in terms of a “nearest-neighbor” thermodynamic model [47–50]. Previous attempts [51] roughly recalibrated only purine/purine and pyrimidine/pyrimidine based on limited experimental data from dinucleotide stacking measurements, which also present challenges in geometrically defining stacking in the absence of the double helix [52]. Our base stacking results provide accurate thermodynamic measurements of single-base stacking for all possible nucleobase combinations in the context of a double helix, which uniquely enables direct calibration of MD force fields in a sequence-specific manner. Here we used our data to evaluate DNA base stacking for two MD force fields optimized for nucleic acids – Amber-99 Chen-Garcia [51] and parmbsc1 [53], the former of which was optimized for RNA, and the latter of which is currently considered the standard force field for MD simulations of DNA (Fig. 6l). To mimic CFM experiments, we simulated two 3-mer duplexes in a solution of ~66000 water molecules and 8 K^+^ ions, enclosed in a 10 nm × 20 nm × 10 nm 3D periodic box. Duplexes were stacked end-to-end and pulled apart (Fig. 6m), with potentials computed as a function of the distance between pulling groups to determine the change in energy between the stacked and unstacked configurations (Fig. 6n). We tested two stacking interactions with pairs of either A|A & T|T or A|T & A|T (Fig. 6o) and found that the parmbsc1 force field overestimated both stacking interactions by ~30-50% while the Amber 99 Chen-Garcia force field underestimated the A|A & T|T and overestimated the A|T & A|T, but was generally closer to experimental values (Fig. 6p). This work shows that our CFM experimental design can be reliably replicated in a MD simulation and used to evaluate and potentially optimize force field parameters to improve the quantitative accuracy of MD simulations for nucleic acids.

## Discussion

This work provides some of the most direct and comprehensive data on base stacking in nucleic acids, while also demonstrating the utility of such detailed knowledge. By employing high-throughput single-molecule experimentation using the CFM combined with novel design of DNA tethers, we measured tens of thousands of individual iterations and quantified base stacking with an uncertainty of ~0.1 kcal/mol. With such small energies, measuring kinetic rates provides an inherent advantage due to the logarithmic dependence of the energies on kinetics. Single molecule techniques are a good fit for this, except they typically make just one measurement at a time. The CFM was developed to address limitations of throughput and accessibility in single-molecule research, and this work marks a milestone as the first large study using the CFM. The throughput and accessibility are evidenced by the ~30,000 single-molecule tethers used in this work, with data collected largely by an undergraduate researcher using a benchtop centrifuge.

Our work provides important new data on base stacking, which generally suggest that previous work has underestimated base stacking energies. One striking example is our measurement of −2.3 kcal/mol for a single G|A stack, which is substantially more stable than measured dinucleotide stacks containing both G|A and T|C, which were reported in the −1.0 to −1.6 kcal/mol range [20,22]. It is likely that a mix of different experimental conditions and biases in experimental designs are responsible for these differences. Our experimental approach provides a fairly direct measurement compared to some other approaches, which included extrapolating stacked/unstacked equilibrium from migration of nicked DNA in urea gels [20], and measuring single-molecule kinetics of on and off rates in end-stacking of DNA origami tubes [22]. One recent paper published during this work used a similar construct design and found a single A|G base stack energy of −2 kcal/mol [54], consistent within error to our measurement. When we compared pairs of our measured base stacking values with previously measured dinucleotide stacks, our energies were larger in all cases by multiples ranging from 1.2 to 2.2.

The data presented here will provide new insights into biological processes, inform DNA design in biotechnology, and improve accuracy for molecular modeling. Especially for short sticky ends that are ubiquitous in biotechnology, base stacking can play a surprisingly large role in stability. Our experimental examples of constructing DNA tetrahedra and monitoring DNA ligation provide glimpses of how our data can be used to tune DNA interactions. While our data was mostly limited to DNA base stacking, our approach can be useful for studying RNA and RNA modifications as well. Our data suggests that RNA base stacking may not be appreciably different from DNA, but further work can help clarify the role of different chemical modifications on base stacking. Our general approach can be adapted to study many variations of nucleotide interactions including those of intercalators under a variety of biologically relevant conditions.

## Acknowledgements

The authors thank Andreas Karl from Thermo Fisher Scientific for providing the centrifuge main board allowing computer control. We thank Pan T.X. Li and Lifeng Zhou for providing feedback on the project. Research reported in this publication was supported by the National Institutes of Health through the National Institute of General Medical Sciences under award R35GM124720 to KH and R35GM133469 to AC, and by the National Science Foundation under award MCB1651877 to AC.

## Author Contributions

The project was conceived and planned by KH and JAP, and supervised by KH. Single-molecule experiments were planned by JAP and were carried out by JAP, KT, AH, and TB. Single-molecule data analysis was performed by JAP and KT. DNA tetrahedra experiments were planned and carried out by ARC. DNA ligation experiments were planned by JAP and carried out by JAP and JV. Molecular dynamics simulations and analysis were planned and carried out by KK, SV, and AC. Technical support and instrument repairs were provided by AH. The paper was written by JAP, ARC, and KH, with MD content by KK, SV, and AC and with general input and editing from other authors.

## References

[1] Li, P. T., Bustamante, C., & Tinoco, I. (2006). Unusual mechanical stability of a minimal RNA kissing complex. Proceedings of the National Academy of Sciences, 103(43), 15847–15852.

[2] Chen, A. A., & García, A. E. (2012). Mechanism of enhanced mechanical stability of a minimal RNA kissing complex elucidated by nonequilibrium molecular dynamics simulations. Proceedings of the National Academy of Sciences, 109(24), E1530–E1539.

[3] Stephenson, W., Asare-Okai, P. N., Chen, A. A., Keller, S., Santiago, R., Tenenbaum, S. A., … & Li, P. T. (2013). The essential role of stacking adenines in a two-base-pair RNA kissing complex. Journal of the American Chemical Society, 135(15), 5602–5611.

[4] Kool, E. T. (2001). Hydrogen bonding, base stacking, and steric effects in DNA replication. Annual review of biophysics and biomolecular structure, 30(1), 1–22.

[5] Reineks, E. Z., & Berdis, A. J. (2004). Evaluating the contribution of base stacking during translesion DNA replication. Biochemistry, 43(2), 393–404.

[6] Takahashi, S., Okura, H., Chilka, P., Ghosh, S., & Sugimoto, N. (2020). Molecular crowding induces primer extension by RNA polymerase through base stacking beyond Watson–Crick rules. RSC Advances, 10(55), 33052–33058.

[7] Lech, C. J., Heddi, B., & Phan, A. T. (2013). Guanine base stacking in G-quadruplex nucleic acids. Nucleic acids research, 41(3), 2034–2046.

[8] Abraham Punnoose, J., Ma, Y., Hoque, M. E., Cui, Y., Sasaki, S., Guo, A. H., … & Mao, H. (2018). Random formation of G-quadruplexes in the full-length human telomere overhangs leads to a kinetic folding pattern with targetable vacant G-tracts. Biochemistry, 57(51), 6946–6955.

[9] Chaires, J. B. (2006). A thermodynamic signature for drug–DNA binding mode. Archives of biochemistry and biophysics, 453(1), 26–31.

[10] Guan, L., & Disney, M. D. (2012). Recent advances in developing small molecules targeting RNA. ACS chemical biology, 7(1), 73–86.

[11] Childs-Disney, J. L., Stepniak-Konieczna, E., Tran, T., Yildirim, I., Park, H., Chen, C. Z., … & Disney, M. D. (2013). Induction and reversal of myotonic dystrophy type 1 pre-mRNA splicing defects by small molecules. Nature communications, 4(1), 1–11.

[12] You, Y., Moreira, B. G., Behlke, M. A., & Owczarzy, R. (2006). Design of LNA probes that improve mismatch discrimination. Nucleic acids research, 34(8), e60–e60.

[13] Loakes, D. (2001). Survey and summary: The applications of universal DNA base analogues. Nucleic Acids Research, 29(12), 2437–2447.

[14] Liu, H., Gao, J., Lynch, S. R., Saito, Y. D., Maynard, L., & Kool, E. T. (2003). A four-base paired genetic helix with expanded size. Science, 302(5646), 868–871.

[15] He, Y., Ye, T., Su, M., Zhang, C., Ribbe, A. E., Jiang, W., & Mao, C. (2008). Hierarchical self-assembly of DNA into symmetric supramolecular 14olyhedral. Nature, 452(7184), 198–201.

[16] Ohayon, Y. P., Hernandez, C., Chandrasekaran, A. R., Wang, X., Abdallah, H. O., Jong, M. A., … & Seeman, N. C. (2019). Designing higher resolution self-assembled 3D DNA crystals via strand terminus modifications. Acs Nano, 13(7), 7957–7965.

[17] Nakata, M., Zanchetta, G., Chapman, B. D., Jones, C. D., Cross, J. O., Pindak, R., … & Clark, N. A. (2007). End-to-end stacking and liquid crystal condensation of 6–to 20–base pair DNA duplexes. Science, 318(5854), 1276–1279.

[18] Wang, R., Kuzuya, A., Liu, W., & Seeman, N. C. (2010). Blunt-ended DNA stacking interactions in a 3-helix motif. Chemical communications, 46(27), 4905–4907.

[19] Liu, L., Li, Y., Wang, Y., Zheng, J., & Mao, C. (2017). Regulating DNA Self□assembly by DNA–Surface Interactions. ChemBioChem, 18(24), 2404–2407.

[20] Protozanova, E., Yakovchuk, P., & Frank-Kamenetskii, M. D. (2004). Stacked–unstacked equilibrium at the nick site of DNA. Journal of molecular biology, 342(3), 775–785.

[21] Yakovchuk, P., Protozanova, E., & Frank-Kamenetskii, M. D. (2006). Base-stacking and base-pairing contributions into thermal stability of the DNA double helix. Nucleic acids research, 34(2), 564–574.

[22] Kilchherr, F., Wachauf, C., Pelz, B., Rief, M., Zacharias, M., & Dietz, H. (2016). Single-molecule dissection of stacking forces in DNA. Science, 353(6304), aaf5508.

[23] Halvorsen, K., & Wong, W. P. (2010). Massively parallel single-molecule manipulation using centrifugal force. Biophysical journal, 98(11), L53–L55.

[24] Kuruvilla, E., Schuster, G. B., & Hud, N. V. (2013). Enhanced Nonenzymatic Ligation of Homopurine Miniduplexes: Support for Greater Base Stacking in a Pre□RNA World. ChemBioChem, 14(1), 45–48.

[25] Cafferty, B. J., Fialho, D. M., Khanam, J., Krishnamurthy, R., & Hud, N. V. (2016). Spontaneous formation and base pairing of plausible prebiotic nucleotides in water. Nature communications, 7(1), 1–8.

[26] Petersheim, M., & Turner, D. H. (1983). Base-stacking and base-pairing contributions to helix stability: thermodynamics of double-helix formation with CCGG, CCGGp, CCGGAp, ACCGGp, CCGGUp, and ACCGGUp. Biochemistry, 22(2), 256–263.

[27] Bommarito, S., Peyret, N., & Jr, J. S. (2000). Thermodynamic parameters for DNA sequences with dangling ends. Nucleic acids research, 28(9), 1929–1934.

[28] Neuman, K. C., & Nagy, A. (2008). Single-molecule force spectroscopy: optical tweezers, magnetic tweezers and atomic force microscopy. Nature methods, 5(6), 491–505.

[29] Yang, D., Ward, A., Halvorsen, K., & Wong, W. P. (2016). Multiplexed single-molecule force spectroscopy using a centrifuge. Nature communications, 7(1), 1–7.

[30] Hoang, T., Patel, D. S., & Halvorsen, K. (2016). A wireless centrifuge force microscope (CFM) enables multiplexed single-molecule experiments in a commercial centrifuge. Review of Scientific Instruments, 87(8), 083705.

[31] Punnoose, J. A., Hayden, A., Zhou, L., & Halvorsen, K. (2020). Wi-Fi Live-Streaming Centrifuge Force Microscope for Benchtop Single-Molecule Experiments. Biophysical journal, 119(11), 2231–2239.

[32] Kirkness, M. W., & Forde, N. R. (2018). Single-molecule assay for proteolytic susceptibility: force-induced collagen destabilization. Biophysical journal, 114(3), 570–576.

[33] LeFevre, T. B., Bikos, D. A., Chang, C. B., & Wilking, J. N. (2021). Measuring colloid–surface interaction forces in parallel using fluorescence centrifuge force microscopy. Soft Matter, 17(26), 6326–6336.

[34] Drew, H. R., Wing, R. M., Takano, T., Broka, C., Tanaka, S., Itakura, K., & Dickerson, R. E. (1981). Structure of a B-DNA dodecamer: conformation and dynamics. Proceedings of the National Academy of Sciences, 78(4), 2179–2183.

[35] Halvorsen, K., Schaak, D., & Wong, W. P. (2011). Nanoengineering a single-molecule mechanical switch using DNA self-assembly. Nanotechnology, 22(49), 494005.

[36] Koussa, M. A., Halvorsen, K., Ward, A., & Wong, W. P. (2015). DNA nanoswitches: a quantitative platform for gel-based biomolecular interaction analysis. Nature methods, 12(2), 123–126.

[37] Bell, G. I. (1978). Models for the specific adhesion of cells to cells: a theoretical framework for adhesion mediated by reversible bonds between cell surface molecules. Science, 200(4342), 618–627.

[38] Evans, E., & Ritchie, K. (1997). Dynamic strength of molecular adhesion bonds. Biophysical journal, 72(4), 1541–1555.

[39] Harcourt, E. M., Kietrys, A. M., & Kool, E. T. (2017). Chemical and structural effects of base modifications in messenger RNA. Nature, 541(7637), 339–346.

[40] Seeman, N. C., & Sleiman, H. F. (2017). DNA nanotechnology. Nature Reviews Materials, 3(1), 1–23.

[41] Hu, Q., Li, H., Wang, L., Gu, H., & Fan, C. (2018). DNA nanotechnology-enabled drug delivery systems. Chemical reviews, 119(10), 6459–6506.

[42] Xiao, M., Lai, W., Man, T., Chang, B., Li, L., Chandrasekaran, A. R., & Pei, H. (2019). Rationally engineered nucleic acid architectures for biosensing applications. Chemical reviews, 119(22), 11631–11717.

[43] Chandrasekaran, A.R. & Halvorsen, K. (2019). Controlled disassembly of a DNA tetrahedron using strand displacement. Nanoscale advances, 1(3), 969–972.

[44] Shuman, S. (2009). DNA ligases: progress and prospects. Journal of Biological Chemistry, 284(26), 17365–17369.

[45] Tomkinson, A. E., Vijayakumar, S., Pascal, J. M., & Ellenberger, T. (2006). DNA ligases: structure, reaction mechanism, and function. Chemical reviews, 106(2), 687–699.

[46] Karplus, M., & Petsko, G. A. (1990). Molecular dynamics simulations in biology. Nature, 347(6294), 631–639.

[47] Tinoco, I., Borer, P. N., Dengler, B., Levine, M. D., Uhlenbeck, O. C., Crothers, D. M., & Gralla, J. (1973). Improved estimation of secondary structure in ribonucleic acids. Nature New Biology, 246(150), 40–41.

[48] Xia, T., SantaLucia Jr, J., Burkard, M. E., Kierzek, R., Schroeder, S. J., Jiao, X., … & Turner, D. H. (1998). Thermodynamic parameters for an expanded nearest-neighbor model for formation of RNA duplexes with Watson− Crick base pairs. Biochemistry, 37(42), 14719–14735.

[49] Lu, Z. J., Turner, D. H., & Mathews, D. H. (2006). A set of nearest neighbor parameters for predicting the enthalpy change of RNA secondary structure formation. Nucleic acids research, 34(17), 4912–4924.

[50] SantaLucia Jr, J. (1998). A unified view of polymer, dumbbell, and oligonucleotide DNA nearest-neighbor thermodynamics. Proceedings of the National Academy of Sciences, 95(4), 1460–1465.

[51] Chen, A. A., & García, A. E. (2013). High-resolution reversible folding of hyperstable RNA tetraloops using molecular dynamics simulations. Proceedings of the National Academy of Sciences, 110(42), 16820–16825.

[52] Taghavi, A., Riveros, I., Wales, D. J., & Yildirim, I. (2022). Evaluating Geometric Definitions of Stacking for RNA Dinucleoside Monophosphates Using Molecular Mechanics Calculations. Journal of Chemical Theory and Computation.

[53] Ivani, I., Dans, P. D., Noy, A., Pérez, A., Faustino, I., Hospital, A., … & Orozco, M. (2016). Parmbsc1: a refined force field for DNA simulations. Nature methods, 13(1), 55–58.

[54] Rieu, M., Vieille, T., Radou, G., Jeanneret, R., Ruiz-Gutierrez, N., Ducos, B., … & Croquette, V. (2021). Parallel, linear, and subnanometric 3D tracking of microparticles with Stereo Darkfield Interferometry. Science Advances, 7(6), eabe3902.

[55] Chandrasekaran, A. R., Dey, B. K., & Halvorsen, K. (2020). How to Perform miRacles: A Step□by□Step microRNA Detection Protocol Using DNA Nanoswitches. Current protocols in molecular biology, 130(1), e114.

[56] Whitley, K. D., Comstock, M. J., & Chemla, Y. R. (2017). Elasticity of the transition state for oligonucleotide hybridization. Nucleic acids research, 45(2), 547–555.

[57] Computing Group ULC; Montreal, QC, Canada: 2021. [(accessed on 29 July 2021)]. Molecular Operating Environment (MOE) Chemical. Available online: https://www.chemcomp.com/Products.htm

[58] Lawrence, C. P., & Skinner, J. L. (2003). Flexible TIP4P model for molecular dynamics simulation of liquid water. Chemical physics letters, 372(5-6), 842–847.

[59] Hess, B., Bekker, H., Berendsen, H. J., & Fraaije, J. G. (1997). LINCS: a linear constraint solver for molecular simulations. Journal of computational chemistry, 18(12), 1463–1472.

[60] Darden, T., York, D., & Pedersen, L. (1993). Particle mesh Ewald: An N⋅ log (N) method for Ewald sums in large systems. The Journal of chemical physics, 98(12), 10089–10092.

[61] Bussi, G., Donadio, D., & Parrinello, M. (2007). Canonical sampling through velocity rescaling. The Journal of chemical physics, 126(1), 014101.

[62] Berendsen, H. J., Postma, J. V., Van Gunsteren, W. F., DiNola, A. R. H. J., & Haak, J. R. (1984). Molecular dynamics with coupling to an external bath. The Journal of chemical physics, 81(8), 3684–3690.

[63] Abraham, M. J., Murtola, T., Schulz, R., Páll, S., Smith, J. C., Hess, B., & Lindahl, E. (2015). GROMACS: High performance molecular simulations through multi-level parallelism from laptops to supercomputers. SoftwareX, 1, 19–25.

