## Supplemental information for "High-throughput single-molecule quantification of individual base stacking energies in nucleic acids"

**Contents**

**Materials and Methods**

**Table S1:** List of oligonucleotides

**Table S2:** Oligonucleotide combinations for each single-molecule construct

**Table S3**: Construct combination to form tethers with preferred base-stacking combination

**Figure S1:** Modular construct design

**Figure S2**: Decay plot and single-exponential fitting of A|C construct at various forces

**Figure S3**: Decay plot and single-exponential fitting of A|T construct at various forces

**Figure S4:** Decay plot and single-exponential fitting of control construct at various forces

**Figure S5:** ΔG_stack_ determined for A|C and A|T base-stack at various forces and at thermal equilibrium

**Figure S6**: Tethering combinations for all combinations

**Figure S7**: Decay plot and single-exponential fitting of A|T, G|T, A|C, and G|C combinations of base-stacks at 15 pN

**Figure S8**: Decay plot and single-exponential fitting of C|T, T|T, C|C combinations and control construct at 15 pN

**Figure S9:** Decay plot and single-exponential fitting of G|A, A|A, G|G combinations of base-stacks corresponding control constructs at 15 pN

**Figure S10**: Tethering combinations for modified base constructs

**Figure S11**: Decay plot and single-exponential fitting of phosphorylated and methylated A|C base-stacks and their corresponding control constructs at 15 pN

**Figure S12**: Decay plot and single-exponential fitting of FAM and ribose A|C base-stacks and their corresponding control constructs at 15 pN

**Figure S13**: Design of DNA tetrahedron.

**Figure S14**: Triplicate gel pics for DNA tetrahedron at various temperatures

**Figure S15:** Triplicate gel pics for ligation experiments with 3 nt overhang

**Figure S16:** Triplicate gel pics for ligation experiments with 4 nt overhang

**Figure S17**: Ligation experiments with 3 nt overhang with G|A Stack vs 4 nt overhang with C|T stack

**Materials and Methods**

*Instrumentation*

The constant force single-molecule experiments in this study were performed using a custom-built CFM, the details of which were largely reported in a previous study [31]. Briefly, the CFM consist of optics comprising a miniaturized video microscope, and of electronics allowing operation and data transmission, in an assembly that fits within a 400 mL bucket of a Sorvall X1R centrifuge. The optical components consist of a 40x plan achromatic infinity-corrected objective (Olympus) for microsphere magnification, turning mirrors (Thorlabs) for achieving required path-length and an LED with diffuser as a light source. The electronic components consist of a gigabit Ethernet machine vision camera (FLIR Blackfly Model # BFLY-PGE-50H5M-C) for imaging, a Wi-Fi router (TP-link TL-WR902AC) for wireless data transfer and communication, and a rechargeable lithium-ion battery (Adafruit) with 5V and 12V voltage step-up regulators (Pololu). These components were assembled within a 3D printed housing (Ultimaker 3). The CFM module and the centrifuge were controlled using a custom written LabVIEW program.

*Sample preparation*

DNA constructs were prepared by hybridizing 124 oligonucleotides (Integrated DNA Technologies) to 7249 nt single-stranded M13mp18 DNA (New England Biolabs). The construction method largely follows our approach for DNA nanoswitch construction [55]. Briefly, the M13 DNA is enzymatically linearized and then incubated with a 10 fold molar excess of backbone oligos 1-122, 150 fold overhang oligo and 500 fold stacking end oligo (Table S2) with an annealing temperature ramp from 90 ˚C to 20 ˚C. In this design, the oligo hybridized to the 3' end of the M13 DNA contains a double biotin on its 5' end for immobilization to streptavidin coated glass surface or the bead. The oligo hybridized to the 5' end of the M13 DNA extends beyond the M13, and provide a platform to anneal an oligo resulting in a 5' single-stranded overhang (Fig. S1). This overhang is used to form ‘sticky-ends’ for different constructs with various base stacking combinations. The list of all oligos used is given in Table S1 and combination of oligos to make constructs with different terminal bases are given in Table S2.

To immobilize DNA constructs to streptavidin coated microspheres (Thermo Fisher Dynabeads M-270 2.8 μm diameter, catalogue # 65306), we used 20 μl of streptavidin microspheres and washed thrice with 50 μl of phosphate-buffered saline containing 0.1% Tween 20 (PBST). Following the washes, the beads solution was brought to a 10 μl volume, and 10 μl of the DNA construct (~500 pM) was added to it and shaken in a vortexer at 1,400 rpm for 30 minutes. The unbound DNA and excess oligos from the construct synthesis was removed by washing the beads thrice with 50 μl PBST and resuspending in 40 μl volume.

The reaction chamber was prepared according to previous work [31]. Briefly, the reaction chamber consists of an 18 mm and a 12 mm circular microscope glass slide (Electron Microscopy Sciences, catalogue # 72230-01 & 72222-01) sandwiching two parallel strips of Kapton tape (www.kaptontape.com) creating a channel of ~ 2 mm between the glass-slides. The glass chamber is assembled on top of a SM1A6 threaded adaptor (Thorlabs). Streptavidin (Amresco) was passively adsorbed to the surface by passing 5 μl of streptavidin (0.1 mg/ml) in 1× PBS. After one minute of incubation, the chamber was washed thrice with 50 μl of PBST to remove unbound streptavidin. Next, 5 μl of DNA construct was passed through the channel and incubated for 10 minutes for the biotin-labeled DNA constructs to bind the streptavidin on the glass surface. The chamber was then washed with PBST to remove unbound constructs and excess oligos from the construct synthesis. DNA coated microspheres were passed into the chamber and incubated for 10 minutes to allow hybridization. The chamber was sealed with vacuum grease and then screwed into the CFM optical assembly until the beads are in focus.

*Constant force experiment protocol*

The prepared CFM with sample chamber was then loaded into the centrifuge bucket, opposite of a counterbalance with matched mass and center of mass. A custom LabView program was used to control the instrument, including the centrifuge speed, image acquisition rate, and camera parameters such as exposure time. The force generated on the tether is the centrifugal force experienced by the beads F = mω^2^r, where m is the effective mass of the bead (actual mass minus the mass of buffer displaced), ω is the angular velocity and r is the distance from the center of the rotor to the chamber (measured at 0.133 m here). The effective mass of beads was determined to be 6.9*10^-12^g for the Dynabeads™ M-270 (www.thermofisher.com) by previous report [23]. The RPM used were 1410, 1221, 997, and 705 for 20pn, 15pN, 10pN, and 5 pN respectively. Experiments were run at a constant force (RPM) for times up to 2 hours and data was saved as individual lossless images.

*Data analysis*

Force induced dissociation of the DNA tethers were measured using a previously reported MATLAB program [31]. The MATLAB program identifies beads using the “imfindcircles” algorithm with a user override for non-spherical, clustered beads and dirt wrongly identified as beads. Once beads are identified from an image at the start of the experiment, the software calculates the variance of the image intensity at the bead location for all the frames. When beads dissociate, it is indicated by the sharp drop in variance (i.e. high contrast to low contrast). Multiple drops in variance observed are due to break in multiple-tethered beads, which are excluded from the analysis. The decay rates were plotted in OriginLab and data was fit using single exponential decay function, y = y_0_ + A×e^-^*^k^*^t^, where y is the fraction of tethers remaining at a given time *t*, y_0_ is the y-axis offset or the baseline, A is the fraction of tethers at the beginning of the experiment (typically 1) and *k* is the off-rate for that particular force. Off-rate for any given condition was determined by at least triplicate experiments where individual k values were determined separately for each set of experiment, and data is reported as the mean and standard deviation of the replicates. The base stacking energies were extracted by comparing the off-rates, given by the equation:

$k_{off}\propto e^{{-E_{a}}/{RT}}$ …………………………………………....……(1)

Where, *E_a_* is the activation energy, *R* is the gas constant and *T* is the absolute temperature. The off-rates of construct with a particular base-stack can be compared to its control construct without base stack to obtain the difference in activation energy:

$\frac{k_{off1}}{k_{off2}}= e^{{{(E}_{a2}-E_{a1})}/{RT}}$ ………………………………..…………(2)

The difference in activation energy between the two constructs is the energy contribution from the base stack following the assumption that the on-rates are equal between the different constructs [56], which can be isolated using the equation:

$\Delta E_{a}=\Delta G_{Base-stack}=RT\ln\left( \frac{k_{off1}}{k_{off2}} \right)$…………………………..(3)

where *k*_off1_ and *k*_off2_ are the off-rates of construct with and without the base stack, *E_a1_* and *E_a2_* are the activation energy barriers for those constructs respectively, and$\Delta G_{Base-stack}$ is the stacking energy of the interfacial bases in the non control construct.

*Assembly and measurement of DNA tetrahedra*

DNA tetrahedra were prepared using previously reported methods [43]. Briefly, DNA strands L, M and S (sequence are shown in Table S1) were mixed in 1:3:3 ratio at 30 nM in Tris-Acetic-EDTA-Mg^2+^ (TAE/Mg^2+^) buffer, which contained 40 mM Tris base (pH 8.0), 20 mM acetic acid, 2 mM EDTA, and 12.5 mM magnesium acetate. The DNA solution was slowly cooled down from 95°C to room temperature over 48 hours in a water bath placed in a Styrofoam box. To assemble DNA tetrahedra with different base stacks, the following strand combinations were used:

1) Tetrahedron with both AG and AT base stacks (control): Strands L, M1, S1
2) Tetrahedron with AG base stack only: Strands L, M2, S1
3) Tetrahedron with AT base stack only: Strands L, M1, S2
4) Tetrahedron with no base stacks: Strands L, M2, S2

DNA tetrahedron assembly was validated using non-denaturing polyacrylamide gel electrophoresis. For gel analysis, 10 μl of the annealed DNA tetrahedron solution was mixed with 1 μl loading dye (containing 50% glycerol and bromophenol blue). 10 μl of this sample was loaded in each gel lane. Gels containing 4% polyacrylamide (29:1 acrylamide/bisacrylamide) were run at 4°C (100 V, constant voltage) in 1X TAE/Mg^2+^ running buffer. After electrophoresis, the gels were stained with GelRed (Biotium) and imaged using Bio-Rad Gel Doc XR+. Gel bands were quantified using ImageJ. To analyze the thermal stability of DNA tetrahedra with different base stacking combinations, we incubated the DNA tetrahedra at 30 °C and 40 °C for 1 hour. Incubated samples were prepared for gel analysis as describe above and tested using 4% PAGE. We quantified the band corresponding to the DNA tetrahedron to obtain the normalized stability levels.

*DNA ligation experiments*

Short duplexes with 20 and 30 bp with 3 or 4 nucleotide overhang were prepared by mixing 50µM oligos and annealing them using temperature ramp from 90 ˚C to 20 ˚C with a temperature gradiant of 1°C/min in 1X PBS buffer (see table S1). To measure the kinetics, 20 and 30 bp duplexes were mixed in equimolar ratio (0.5µM) in buffer with final concentration of 1X T4 DNA ligase buffer (NEB), 1X BSA, 1 mM ATP. 1 µl T4 DNA ligase (40 units/ µl) was added to the mixture. The reaction was terminated at the required time point by heat inactivation at 70°C for 20 mins. Then the reaction mixtures were mixed with the gel loading buffer (final concentration 1X) and were run in a 10% non-denaturing PAGE (29:1 acrylamide/bisacrylamide) at room temperature (150 V, 1hr). After electrophoresis, the gels were stained with GelRed (Biotium) and imaged using Bio-Rad Gel Doc XR+.Gel bands were quantified using ImageLabs software. The ligated product was quantified and normalized against the 50 bp marker band in the 10 bp-DNA ladder (Thermofisher, Catalog number: SM1313)

*Molecular dynamics simulations*

Two end-to-end stacked 3-mer duplexes were pulled apart using molecular dynamics (MD) simulations to mimic CFM experiments. We evaluated the stacking parameters of two nucleic acid force fields:  Amber-99 with Chen-Garcia correction [51] and parmbsc1 [53]. The initial structures were constructed using the Molecular Operating Environment software [57] and consist of sequences 5’-CGX|xTC-3’ and 3’-GCY|yAG-5’, where | indicates the boundary of each duplex; Xx-Yy are either AA-TT, or AT-TA, respectively. The simulation system consisted of the two 3-mer DNA duplexes in a solution of ~66000 water molecules and 8 K+ ions, enclosed in a 10nm × 20nm × 10nm 3D periodic box. Water molecules were represented with the TIP4P model [58] and LINCS was used to restrain hydrogens bonded to heavy atoms [59].  Long-ranged electrostatic interactions were calculated using particle mesh Ewald (PME) algorithm [60]. The system was subjected to steepest-descent energy minimization, followed by NVT and NPT equilibration runs, 1 ns each maintained at 310 K using the velocity rescaling thermostat [61] and at 1 atm using the Berendsen barostat [62]. Sixteen parallel MD pulling simulations were then performed for 2.6—2.8 ns /simulation, with varying initial distances (0.475nm & 0.5-1.2 nm in 0.05nm intervals) between the pull groups (C1’ atoms on one of the stacked base pair), totaling ~42 ns of production run per construct per forcefield. The simulations incorporated leap-frog algorithm with a 2 fs time-step in the NVT ensemble at 310 K using the velocity rescaling thermostat [61]. The base-pairing between the two base pairs closest to the 3’ and 5’ ends was maintained using distance restraints with a force constant of 1195 kcal·mol^-1^·nm^-2^. A pulling force constant 1600 kcal·mol^-1^·nm^-2^ was used for all cases except the AA-TT: bsc1 simulation which required a higher force (8000 kcal·mol^-1^·nm^-2^) to pull apart the duplexes within the timeframe of the simulation. System coordinates were stored every 2 ps. All MD simulations were performed using Gromacs-2020.01 package and the potential mean force (PMF) was extracted using the Weighted Histogram Analysis Method (WHAM) module [63].

**Table S1.** List of oligonucleotide sequences.

| **Name** | **Sequence** | **Length** | |
| --- | --- | --- | --- |
| **Backbone sequences (5′-3′)**  **(Common set of oligos for all single-molecule constructs)** | | | |
| 1. 5’Biotin | (5’ 2x bio) AACATCCAATAAATCATACAGGCAAGGCAAAGAATTAGCA | 40 | |
| 2 | AAATTAAGCAATAAAGCCTC | 20 | |
| 3 | AGAGCATAAAGCTAAATCGGTTGTACCAAAAACATTATGACCCTGTAATACTTTTGCGGG | 60 | |
| 4 | AGAAGCCTTTATTTCAACGCAAGGATAAAAATTTTTAGAACCCTCATATATTTTAAATGC | 60 | |
| 5 | AATGCCTGAGTAATGTGTAGGTAAAGATTCAAAAGGGTGAGAAAGGCCGGAGACAGTCAA | 60 | |
| 6 | ATCACCATCAATATGATATTCAACCGTTCTAGCTGATAAATTAATGCCGGAGAGGGTAGC | 60 | |
| 7 | TATTTTTGAGAGATCTACAAAGGCTATCAGGTCATTGCCTGAGAGTCTGGAGCAAACAAG | 60 | |
| 8 | AGAATCGATGAACGGTAATCGTAAAACTAGCATGTCAATCATATGTACCCCGGTTGATAA | 60 | |
| 9 | TCAGAAAAGCCCCAAAAACAGGAAGATTGTATAAGCAAATATTTAAATTGTAAACGTTAA | 60 | |
| 10 | TATTTTGTTAAAATTCGCATTAAATTTTTGTTAAATCAGCTCATTTTTTAACCAATAGGA | 60 | |
| 11 | ACGCCATCAAAAATAATTCGCGTCTGGCCTTCCTGTAGCCAGCTTTCATCAACATTAAAT | 60 | |
| 12 | GTGAGCGAGTAACAACCCGTCGGATTCTCCGTGGGAACAAACGGCGGATTGACCGTAATG | 60 | |
| 13 | GGATAGGTCACGTTGGTGTAGATGGGCGCATCGTAACCGTGCATCTGCCAGTTTGAGGGG | 60 | |
| 14 | ACGACGACAGTATCGGCCTCAGGAAGATCGCACTCCAGCCAGCTTTCCGGCACCGCTTCT | 60 | |
| 15 | GGTGCCGGAAACCAGGCAAAGCGCCATTCGCCATTCAGGCTGCGCAACTGTTGGGAAGGG | 60 | |
| 16 | CGATCGGTGCGGGCCTCTTCGCTATTACGCCAGCTGGCGAAAGGGGGATGTGCTGCAAGG | 60 | |
| 17 | CGATTAAGTTGGGTAACGCCAGGGTTTTCCCAGTCACGACGTTGTAAAACGACGGCCAGT | 60 | |
| 18 | GCCAAGCTTGCATGCCTGCAGGTCGACTCTAGAGGATCCCCGGGTACCGAGCTCGAATTC | 60 | |
| 19 | GTAATCATGGTCATAGCTGTTTCCTGTGTGAAATTGTTATCCGCTCACAATTCCACACAA | 60 | |
| 20 | CATACGAGCCGGAAGCATAAAGTGTAAAGCCTGGGGTGCCTAATGAGTGAGCTAACTCAC | 60 | |
| 21 | ATTAATTGCGTTGCGCTCACTGCCCGCTTTCCAGTCGGGAAACCTGTCGTGCCAGCTGCA | 60 | |
| 22 | TTAATGAATCGGCCAACGCGCGGGGAGAGGCGGTTTGCGTATTGGGCGCCAGGGTGGTTT | 60 | |
| 23 | TTCTTTTCACCAGTGAGACGGGCAACAGCTGATTGCCCTTCACCGCCTGGCCCTGAGAGA | 60 | |
| 24 | GTTGCAGCAAGCGGTCCACGCTGGTTTGCCCCAGCAGGCGAAAATCCTGTTTGATGGTGG | 60 | |
| 25 | TTCCGAAATCGGCAAAATCCCTTATAAATCAAAAGAATAGCCCGAGATAGGGTTGAGTGT | 60 | |
| 26 | TGTTCCAGTTTGGAACAAGAGTCCACTATTAAAGAACGTGGACTCCAACGTCAAAGGGCG | 60 | |
| 27 | AAAAACCGTCTATCAGGGCGATGGCCCACTACGTGAACCATCACCCAAATCAAGTTTTTT | 60 | |
| 28 | GGGGTCGAGGTGCCGTAAAGCACTAAATCGGAACCCTAAAGGGAGCCCCCGATTTAGAGC | 60 | |
| 29 | TTGACGGGGAAAGCCGGCGAACGTGGCGAGAAAGGAAGGGAAGAAAGCGAAAGGAGCGGG | 60 | |
| 30 | CGCTAGGGCGCTGGCAAGTGTAGCGGTCACGCTGCGCGTAACCACCACACCCGCCGCGCT | 60 | |
| 31 | TAATGCGCCGCTACAGGGCGCGTACTATGGTTGCTTTGACGAGCACGTATAACGTGCTTT | 60 | |
| 32 | CCTCGTTAGAATCAGAGCGGGAGCTAAACAGGAGGCCGATTAAAGGGATTTTAGACAGGA | 60 | |
| 33 | ACGGTACGCCAGAATCCTGAGAAGTGTTTTTATAATCAGTGAGGCCACCGAGTAAAAGAG | 60 | |
| 34 | TCTGTCCATCACGCAAATTAACCGTTGTAGCAATACTTCTTTGATTAGTAATAACATCAC | 60 | |
| 35 | TTGCCTGAGTAGAAGAACTCAAACTATCGGCCTTGCTGGTAATATCCAGAACAATATTAC | 60 | |
| 36 | CGCCAGCCATTGCAACAGGAAAAACGCTCATGGAAATACCTACATTTTGACGCTCAATCG | 60 | |
| 37 | TCTGAAATGGATTATTTACATTGGCAGATTCACCAGTCACACGACCAGTAATAAAAGGGA | 60 | |
| 38 | CATTCTGGCCAACAGAGATAGAACCCTTCTGACCTGAAAGCGTAAGAATACGTGGCACAG | 60 | |
| 39 | ACAATATTTTTGAATGGCTATTAGTCTTTAATGCGCGAACTGATAGCCCTAAAACATCGC | 60 | |
| 40 | CATTAAAAATACCGAACGAACCACCAGCAGAAGATAAAACAGAGGTGAGGCGGTCAGTAT | 60 | |
| 41 | TAACACCGCCTGCAACAGTGCCACGCTGAGAGCCAGCAGCAAATGAAAAATCTAAAGCAT | 60 | |
| 42 | CACCTTGCTGAACCTCAAATATCAAACCCTCAATCAATATCTGGTCAGTTGGCAAATCAA | 60 | |
| 43 | CAGTTGAAAGGAATTGAGGAAGGTTATCTAAAATATCTTTAGGAGCACTAACAACTAATA | 60 | |
| 44 | GATTAGAGCCGTCAATAGATAATACATTTGAGGATTTAGAAGTATTAGACTTTACAAACA | 60 | |
| 45 | ATTCGACAACTCGTATTAAATCCTTTGCCCGAACGTTATTAATTTTAAAAGTTTGAGTAA | 60 | |
| 46 | CATTATCATTTTGCGGAACAAAGAAACCACCAGAAGGAGCGGAATTATCATCATATTCCT | 60 | |
| 47 | GATTATCAGATGATGGCAATTCATCAATATAATCCTGATTGTTTGGATTATACTTCTGAA | 60 | |
| 48 | TAATGGAAGGGTTAGAACCTACCATATCAAAATTATTTGCACGTAAAACAGAAATAAAGA | 60 | |
| 49 | AATTGCGTAGATTTTCAGGTTTAACGTCAGATGAATATACAGTAACAGTACCTTTTACAT | 60 | |
| 50 | CGGGAGAAACAATAACGGATTCGCCTGATTGCTTTGAATACCAAGTTACAAAATCGCGCA | 60 | |
| 51 | GAGGCGAATTATTCATTTCAATTACCTGAGCAAAAGAAGATGATGAAACAAACATCAAGA | 60 | |
| 52 | AAACAAAATTAATTACATTTAACAATTTCATTTGAATTACCTTTTTTAATGGAAACAGTA | 60 | |
| 53 | CATAAATCAATATATGTGAGTGAATAACCTTGCTTCTGTAAATCGTCGCTATTAATTAAT | 60 | |
| 54 | TTTCCCTTAGAATCCTTGAAAACATAGCGATAGCTTAGATTAAGACGCTGAGAAGAGTCA | 60 | |
| 55 | ATAGTGAATTTATCAAAATCATAGGTCTGAGAGACTACCTTTTTAACCTCCGGCTTAGGT | 60 | |
| 56 | TGGGTTATATAACTATATGTAAATGCTGATGCAAATCCAATCGCAAGACAAAGAACGCGA | 60 | |
| 57 | GAAAACTTTTTCAAATATATTTTAGTTAATTTCATCTTCTGACCTAAATTTAATGGTTTG | 60 | |
| 58 | AAATACCGACCGTGTGATAAATAAGGCGTTAAATAAGAATAAACACCGGAATCATAATTA | 60 | |
| 59 | CTAGAAAAAGCCTGTTTAGTATCATATGCGTTATACAAATTCTTACCAGTATAAAGCCAA | 60 | |
| 60 | CGCTCAACAGTAGGGCTTAATTGAGAATCGCCATATTTAACAACGCCAACATGTAATTTA | 60 | |
| 61 | GGCAGAGGCATTTTCGAGCCAGTAATAAGAGAATATAAAGTACCGACAAAAGGTAAAGTA | 60 | |
| 62 | ATTCTGTCCAGACGACGACAATAAACAACATGTTCAGCTAATGCAGAACGCGCCTGTTTA | 60 | |
| 63 | TCAACAATAGATAAGTCCTGAACAAGAAAAATAATATCCCATCCTAATTTACGAGCATGT | 60 | |
| 64 | AGAAACCAATCAATAATCGGCTGTCTTTCCTTATCATTCCAAGAACGGGTATTAAACCAA | 60 | |
| 65 | GTACCGCACTCATCGAGAACAAGCAAGCCGTTTTTATTTTCATCGTAGGAATCATTACCG | 60 | |
| 66 | CGCCCAATAGCAAGCAAATCAGATATAGAAGGCTTATCCGGTATTCTAAGAACGCGAGGC | 60 | |
| 67 | GTTTTAGCGAACCTCCCGACTTGCGGGAGGTTTTGAAGCCTTAAATCAAGATTAGTTGCT | 60 | |
| 68 | ATTTTGCACCCAGCTACAATTTTATCCTGAATCTTACCAACGCTAACGAGCGTCTTTCCA | 60 | |
| 69 | GAGCCTAATTTGCCAGTTACAAAATAAACAGCCATATTATTTATCCCAATCCAAATAAGA | 60 | |
| 70 | AACGATTTTTTGTTTAACGTCAAAAATGAAAATAGCAGCCTTTACAGAGAGAATAACATA | 60 | |
| 71 | AAAACAGGGAAGCGCATTAGACGGGAGAATTAACTGAACACCCTGAACAAAGTCAGAGGG | 60 | |
| 72 | TAATTGAGCGCTAATATCAGAGAGATAACCCACAAGAATTGAGTTAAGCCCAATAATAAG | 60 | |
| 73 | AGCAAGAAACAATGAAATAGCAATAGCTATCTTACCGAAGCCCTTTTTAAGAAAAGTAAG | 60 | |
| 74 | CAGATAGCCGAACAAAGTTACCAGAAGGAAACCGAGGAAACGCAATAATAACGGAATACC | 60 | |
| 75 | CAAAAGAACTGGCATGATTAAGACTCCTTATTACGCAGTATGTTAGCAAACGTAGAAAAT | 60 | |
| 76 | ACATACATAAAGGTGGCAACATATAAAAGAAACGCAAAGACACCACGGAATAAGTTTATT | 60 | |
| 77 | TTGTCACAATCAATAGAAAATTCATATGGTTTACCAGCGCCAAAGACAAAAGGGCGACAT | 60 | |
| 78 | TCAACCGATTGAGGGAGGGAAGGTAAATATTGACGGAAATTATTCATTAAAGGTGAATTA | 60 | |
| 79 | TCACCGTCACCGACTTGAGCCATTTGGGAATTAGAGCCAGCAAAATCACCAGTAGCACCA | 60 | |
| 80 | TTACCATTAGCAAGGCCGGAAACGTCACCAATGAAACCATCGATAGCAGCACCGTAATCA | 60 | |
| 81 | GTAGCGACAGAATCAAGTTTGCCTTTAGCGTCAGACTGTAGCGCGTTTTCATCGGCATTT | 60 | |
| 82 | TCGGTCATAGCCCCCTTATTAGCGTTTGCCATCTTTTCATAATCAAAATCACCGGAACCA | 60 | |
| 83 | GAGCCACCACCGGAACCGCCTCCCTCAGAGCCGCCACCCTCAGAACCGCCACCCTCAGAG | 60 | |
| 84 | CCACCACCCTCAGAGCCGCCACCAGAACCACCACCAGAGCCGCCGCCAGCATTGACAGGA | 60 | |
| 85 | GGTTGAGGCAGGTCAGACGATTGGCCTTGATATTCACAAACAAATAAATCCTCATTAAAG | 60 | |
| 86 | CCAGAATGGAAAGCGCAGTCTCTGAATTTACCGTTCCAGTAAGCGTCATACATGGCTTTT | 60 | |
| 87 | GATGATACAGGAGTGTACTGGTAATAAGTTTTAACGGGGTCAGTGCCTTGAGTAACAGTG | 60 | |
| 88 | CCCGTATAAACAGTTAATGCCCCCTGCCTATTTCGGAACCTATTATTCTGAAACATGAAA | 60 | |
| 89 | GTATTAAGAGGCTGAGACTCCTCAAGAGAAGGATTAGGATTAGCGGGGTTTTGCTCAGTA | 60 | |
| 90 | CCAGGCGGATAAGTGCCGTCGAGAGGGTTGATATAAGTATAGCCCGGAATAGGTGTATCA | 60 | |
| 91 | CCGTACTCAGGAGGTTTAGTACCGCCACCCTCAGAACCGCCACCCTCAGAACCGCCACCC | 60 | |
| 92 | TCAGAGCCACCACCCTCATTTTCAGGGATAGCAAGCCCAATAGGAACCCATGTACCGTAA | 60 | |
| 93 | CACTGAGTTTCGTCACCAGTACAAACTACAACGCCTGTAGCATTCCACAGACAGCCCTCA | 60 | |
| 94 | TAGTTAGCGTAACGATCTAAAGTTTTGTCGTCTTTCCAGACGTTAGTAAATGAATTTTCT | 60 | |
| 95 | GTATGGGATTTTGCTAAACAACTTTCAACAGTTTCAGCGGAGTGAGAATAGAAAGGAACA | 60 | |
| 96 | ACTAAAGGAATTGCGAATAATAATTTTTTCACGTTGAAAATCTCCAAAAAAAAGGCTCCA | 60 | |
| 97 | AAAGGAGCCTTTAATTGTATCGGTTTATCAGCTTGCTTTCGAGGTGAATTTCTTAAACAG | 60 | |
| 98 | CTTGATACCGATAGTTGCGCCGACAATGACAACAACCATCGCCCACGCATAACCGATATA | 60 | |
| 99 | TTCGGTCGCTGAGGCTTGCAGGGAGTTAAAGGCCGCTTTTGCGGGATCGTCACCCTCAGC | 60 | |
| 100 | AGCGAAAGACAGCATCGGAACGAGGGTAGCAACGGCTACAGAGGCTTTGAGGACTAAAGA | 60 | |
| 101 | CTTTTTCATGAGGAAGTTTCCATTAAACGGGTAAAATACGTAATGCCACTACGAAGGCAC | 60 | |
| 102 | CAACCTAAAACGAAAGAGGCAAAAGAATACACTAAAACACTCATCTTTGACCCCCAGCGA | 60 | |
| 103 | TTATACCAAGCGCGAAACAAAGTACAACGGAGATTTGTATCATCGCCTGATAAATTGTGT | 60 | |
| 104 | CGAAATCCGCGACCTGCTCCATGTTACTTAGCCGGAACGAGGCGCAGACGGTCAATCATA | 60 | |
| 105 | AGGGAACCGAACTGACCAACTTTGAAAGAGGACAGATGAACGGTGTACAGACCAGGCGCA | 60 | |
| 106 | TAGGCTGGCTGACCTTCATCAAGAGTAATCTTGACAAGAACCGGATATTCATTACCCAAA | 60 | |
| 107 | TCAACGTAACAAAGCTGCTCATTCAGTGAATAAGGCTTGCCCTGACGAGAAACACCAGAA | 60 | |
| 108 | CGAGTAGTAAATTGGGCTTGAGATGGTTTAATTTCAACTTTAATCATTGTGAATTACCTT | 60 | |
| 109 | ATGCGATTTTAAGAACTGGCTCATTATACCAGTCAGGACGTTGGGAAGAAAAATCTACGT | 60 | |
| 110 | TAATAAAACGAACTAACGGAACAACATTATTACAGGTAGAAAGATTCATCAGTTGAGATT | 60 | |
| 111 | TAGGAATACCACATTCAACTAATGCAGATACATAACGCCAAAAGGAATTACGAGGCATAG | 60 | |
| 112 | TAAGAGCAACACTATCATAACCCTCGTTTACCAGACGACGATAAAAACCAAAATAGCGAG | 60 | |
| 113 | AGGCTTTTGCAAAAGAAGTTTTGCCAGAGGGGGTAATAGTAAAATGTTTAGACTGGATAG | 60 | |
| 114 | CGTCCAATACTGCGGAATCGTCATAAATATTCATTGAATCCCCCTCAAATGCTTTAAACA | 60 | |
| 115 | GTTCAGAAAACGAGAATGACCATAAATCAAAAATCAGGTCTTTACCCTGACTATTATAGT | 60 | |
| 116 | CAGAAGCAAAGCGGATTGCATCAAAAAGATTAAGAGGAAGCCCGAAAGACTTCAAATATC | 60 | |
| 117 | GCGTTTTAATTCGAGCTTCAAAGCGAACCAGACCGGAAGCAAACTCCAACAGGTCAGGAT | 60 | |
| 118 | TAGAGAGTACCTTTAATTGCTCCTTTTGATAAGAGGTCATTTTTGCGGATGGCTTAGAGC | 60 | |
| 119 | TTAATTGCTGAATATAATGCTGTAGCTCAACATGTTTTAAATATGCAACTAAAGTACGGT | 60 | |
| 120 | GTCTGGAAGTTTCATTCCATATAACAGTTGATTCCCAATTCTGCGAACGAGTAGATTTAG | 60 | |
| 121 | TTTGACCATTAGATACATTTCGCAAATGGTCAATAACCTGTTTAGCTAT | 49 | |
| 122 | ATTTTCATTTGGGGCGCGAGCTGAAAAGGT | 30 | |
| **Sequence used in specific combination for each construct (5′-3′)**  **Overhanging regions underlined, spacer T are marked in blue, and stacking bases in red** | | | |
| OH-A | GGCATCAATTCTACTAATAGTAGTAGCATTCCGTGCCTGTGAACGAGCTGCCCCATGGCA | 60 | |
| OH-G | GGCATCAATTCTACTAATAGTAGTAGCATTCCGTGCCTGTGAACGAGCTGCCCCATGGCG | 60 | |
| OH-C | GGCATCAATTCTACTAATAGTAGTAGCATTCCGTGCCTGTGAACGAGCTGCCCCATGGCC | 60 | |
| OH-T | GGCATCAATTCTACTAATAGTAGTAGCATTCCGTGCCTGTGAACGAGCTGCCCCATGGCT | 60 | |
| A:A-C | CCGCTGCATGCCATGGGGCAGCTCGTTCACAGGCACGG | 38 | |
| G:A-C | CCGCTGCACGCCATGGGGCAGCTCGTTCACAGGCACGG | 38 | |
| C:A-C | CCGCTGCAGGCCATGGGGCAGCTCGTTCACAGGCACGG | 38 | |
| T:A-C | CCGCTGCAAGCCATGGGGCAGCTCGTTCACAGGCACGG | 38 | |
| T: Sp-A-C | CCGCTGCATTTAGCCATGGGGCAGCTCGTTCACAGGCACGG | 41 | |
| A: G-T | TGCAGCGGTGCCATGGGGCAGCTCGTTCACAGGCACGG | 38 | |
| G: G-T | TGCAGCGGCGCCATGGGGCAGCTCGTTCACAGGCACGG | 38 | |
| C: G-T | TGCAGCGGGGCCATGGGGCAGCTCGTTCACAGGCACGG | 38 | |
| T: G-T | TGCAGCGGAGCCATGGGGCAGCTCGTTCACAGGCACGG | 38 | |
| T:Sp-G-T | TGCAGCGGTTTAGCCATGGGGCAGCTCGTTCACAGGCACGG | 41 | |
| G: C-A | ACGTCGCCCGCCATGGGGCAGCTCGTTCACAGGCACGG | 38 | |
| T: Sp-C-A | ACGTCGCCTTTAGCCATGGGGCAGCTCGTTCACAGGCACGG | 41 | |
| A: T-G | GGCGACGTTGCCATGGGGCAGCTCGTTCACAGGCACGG | 38 | |
| G: T-G | GGCGACGTCGCCATGGGGCAGCTCGTTCACAGGCACGG | 38 | |
| T: Sp-T-G | GGCGACGTTTTAGCCATGGGGCAGCTCGTTCACAGGCACGG | 41 | |
| **Sequence used for constructs with modified stacks (5′-3′)**  **Overhanging regions underlined, spacer T are marked in blue, and stacking bases in red** | | | |
| T:Sp-A-C^p^ | /5Phos/CCGCTGCATTTAGCCATGGGGCAGCTCGTTCACAGGCACGG | | 41 |
| T:Sp-A-C^FAM^ | /56-FAM/CCGCTGCATTTAGCCATGGGGCAGCTCGTTCACAGGCACGG | | 41 |
| T:Sp-A-^Me^C | /5Me-dC/CGCTGCATTTAGCCATGGGGCAGCTCGTTCACAGGCACGG | | 41 |
| OH-rA | GGCATCAATTCTACTAATAGTAGTAGCATTCCGTGCCTGTGAACGAGCTGCCCCATGGCrA | | 60 |
| **Sequence used for constructing DNA tetrahedron (5′-3′)** | | | |
| L | AGGCACCATCGTAGGTTTTTCTTGCCAGGCACCATCGTAGGTTTTTCTTGCCAGGCACCATCG TAGGTTTTTCTTGCC | | 78 |
| M1 | TAGCAACCTGCCTGGCAAGCCTACGATGGACACGGTATCGCA | | 42 |
| S1 | ATACCGTGTGGTTGCTATGCG | | 21 |
| M2 | AGCAACCTGCCTGGCAAGCCTACGATGGACACGGTATCGCA | | 41 |
| S2 | TACCGTGTGGTTGCTATGCG | | 20 |
| **Sequences used for Ligation experiments (5′-3′)**  **Overhanging regions underlined and stacking bases are shown in red** | | | |
| 20 AG | TATGTGTCTTCTCAGTAGTG | | 20 |
| 3 + 20 AG | /5Phos/ACTCACTACTGAGAAGACACATA | | 23 |
| 4 + 20 AG | /5Phos/ACCTCACTACTGAGAAGACACATA | | 24 |
| 30 AG | ATACTTATACAGCTTATTGACCATTTGCGG | | 30 |
| 3 + 30 AG | /5Phos/AGTCCGCAAATGGTCAATAAGCTGTATAAGTAT | | 33 |
| 4 + 30 AG | /5Phos/AGGTCCGCAAATGGTCAATAAGCTGTATAAGTAT | | 34 |
| 20 AT | TATGTGTCTTCTCAGTAGTT | | 20 |
| 3 + 20 AT | /5Phos/ACTAACTACTGAGAAGACACATA | | 23 |
| 4 + 20 AT | /5Phos/ACCTAACTACTGAGAAGACACATA | | 24 |
| 30 AT | ATACTTATACAGCTTATTGACCATTTGCGT | | 30 |
| 3 + 30 AT | /5Phos/AGTACGCAAATGGTCAATAAGCTGTATAAGTAT | | 33 |
| 4 + 30 AT | /5Phos/AGGTACGCAAATGGTCAATAAGCTGTATAAGTAT | | 34 |

**Table S2:** Oligonucleotide combinations for each single-molecule construct.

| **Construct #** | **Oligo Mix (1:15:50 mole ratio)** |
| --- | --- |
| 1 | Oligos 1-122, OH-A, A:A-C |
| 2 | Oligos 1-122, OH-G, G:A-C |
| 3 | Oligos 1-122, OH-C, C:A-C |
| 4 | Oligos 1-122, OH-T, T:A-C |
| 5 | Oligos 1-122, OH-T, T:Sp-G-T |
| 6 | Oligos 1-122, OH-A, A:G-T |
| 7 | Oligos 1-122, OH-G, G:G-T |
| 8 | Oligos 1-122, OH-C, C:G-T |
| 9 | Oligos 1-122, OH-T, T:G-T |
| 10 | Oligos 1-122, OH-T, T:Sp-A-C |
| 11 | Oligos 1-122, OH-G, G:T-G |
| 12 | Oligos 1-122, OH-A, A:T-G |
| 13 | Oligos 1-122, OH-T, T:Sp-C-A |
| 14 | Oligos 1-122, OH-G, G:C-A |
| 15 | Oligos 1-122, OH-T, T:Sp-T-G |
| 16 | Oligos 1-122, OH-T, T:Sp-A-C^p^ |
| 17 | Oligos 1-122, OH-T, T:Sp-A-C^me^ |
| 18 | Oligos 1-122, OH-T, T:Sp-A-C^FAM^ |
| 19 | Oligos 1-122, OH-rA, A:T-G |

**Table S3:** Construct combination to form tethers in single-molecule experiments**.**

| **Tether ID** | **Stacking Combination (5′\|3′)** | **Construct combinations to form tether** |
| --- | --- | --- |
| 1 | A\|T | 1 & 5 |
| 2 | G\|T | 2 & 5 |
| 3 | C\|T | 3 & 5 |
| 4 | T\|T | 4 & 5 |
| 5 | A\|C | 6 & 10 |
| 6 | G\|C | 7 & 10 |
| 7 | C\|C | 8 & 10 |
| 8 | Control for (A\|T, G\|T, C\|T, T\|T, A\|C, G\|C, C\|C) | 5 & 10 |
| 9 | G\|A | 11 & 13 |
| 10 | A\|A | 12 & 13 |
| 11 | G\|G | 14 & 15 |
| 12 | Control for (G\|A, A\|A, G\|G) | 13 & 15 |
| 13 | A\| ^p^C | 16 & 6 |
| 14 | Control for A\|^p^C | 16 & 5 |
| 15 | A\| ^me^C | 17 & 6 |
| 16 | Control A\|^me^C | 17 & 5 |
| 17 | A ^FAM^\|C | 18 & 6 |
| 18 | Control A\|^FAM^C | 18 & 5 |
| 19 | rA\|C | 19 & 6 |
| 8 | Control for rA\|C | 5 & 10 |

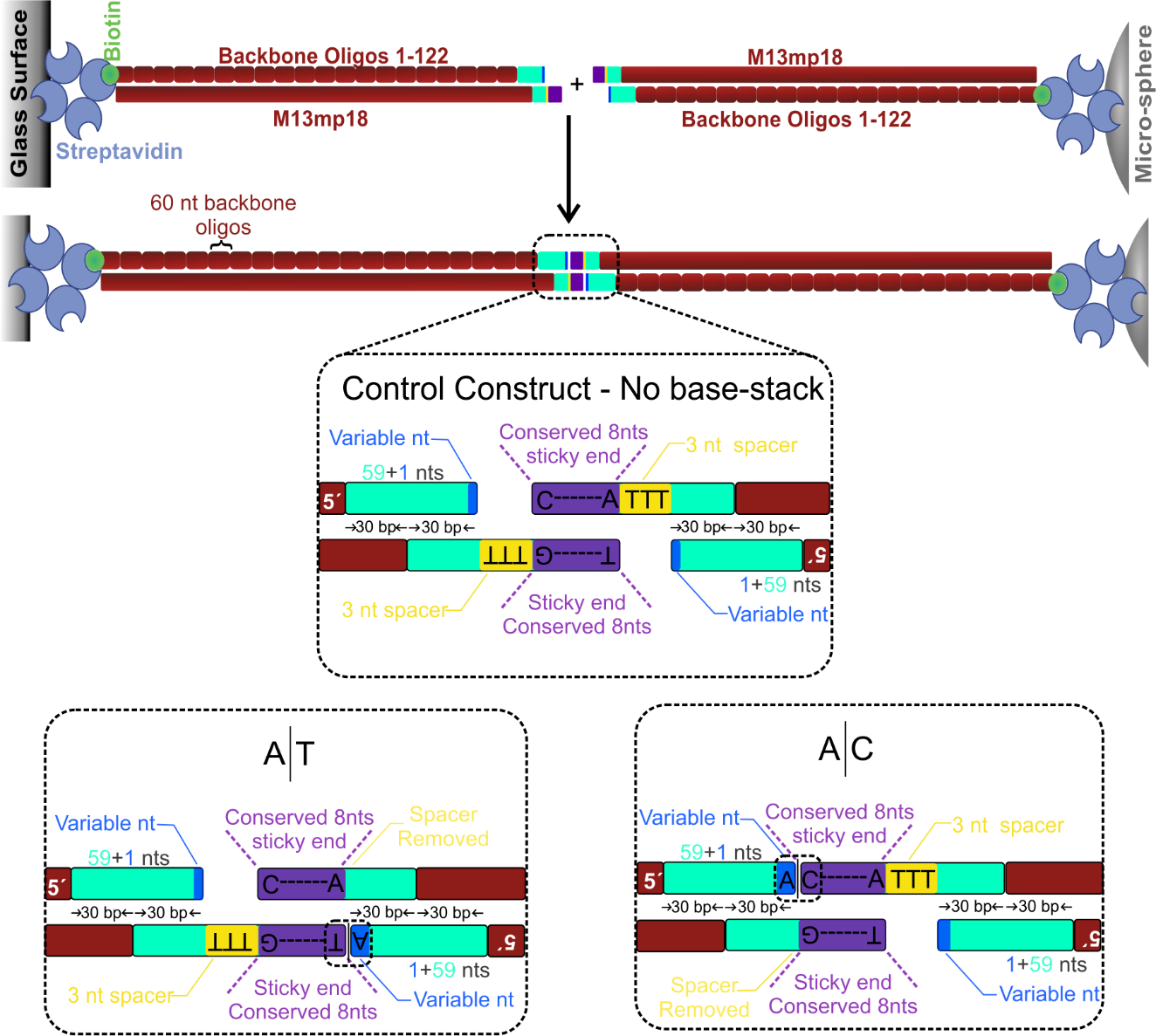

***Figure S1: Modular construct design.*** *The central duplex (purple) has 4 nucleotides in its termini, which can interface with the variable region (blue) to form the base-stack of interest in the absence of 3 nt spacer nucleotides (yellow). The enlarged central region shows the tethers that has an A|T and an A|C base-stack and the control construct without any base-stacks. All these constructs have the same central duplex forming the same base-pairs, thus the difference in the tether strength is due to the base-stack of interest.*

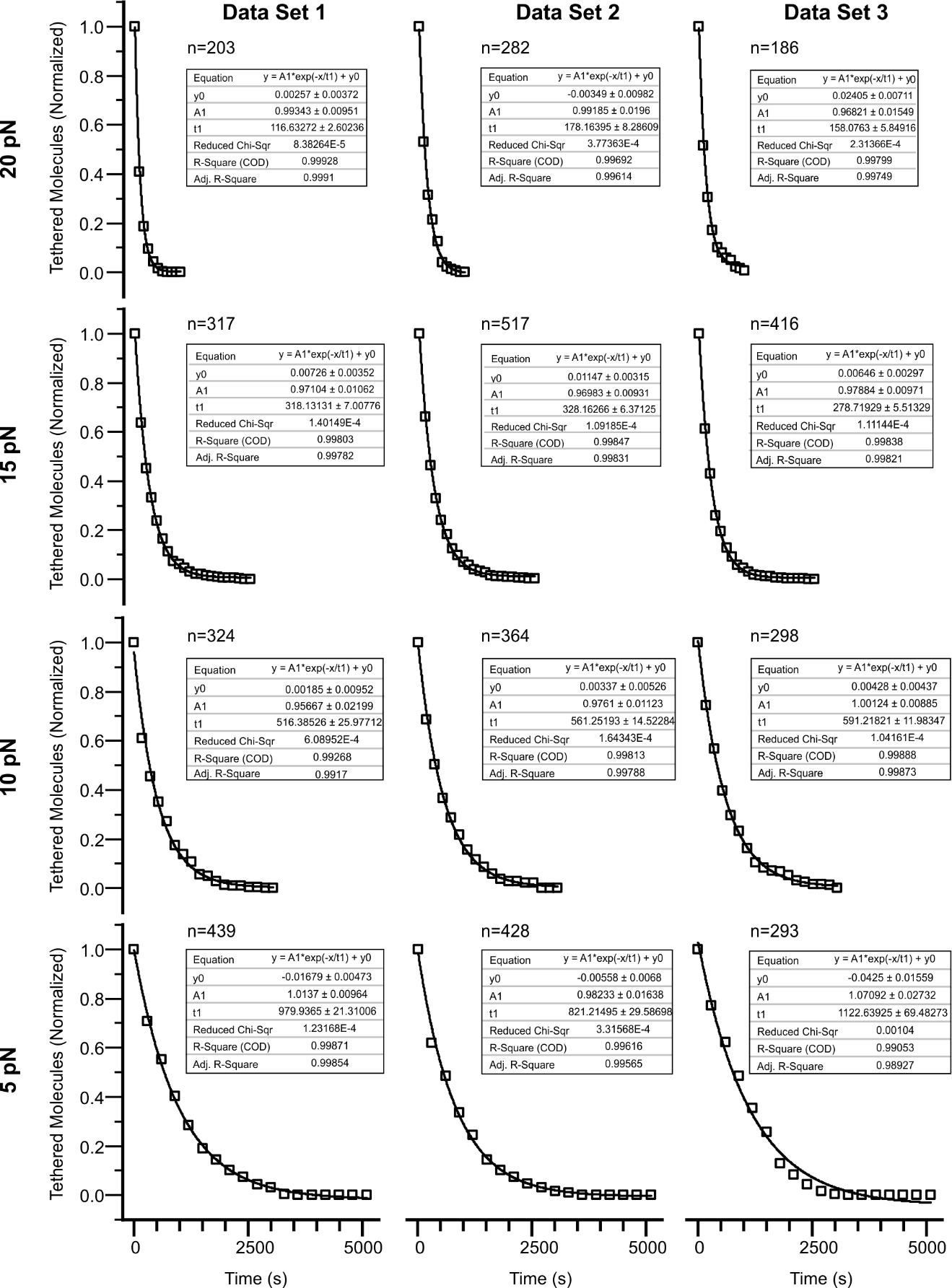

***Figure S2: Decay plot and single-exponential fitting of A|C construct at various forces.***

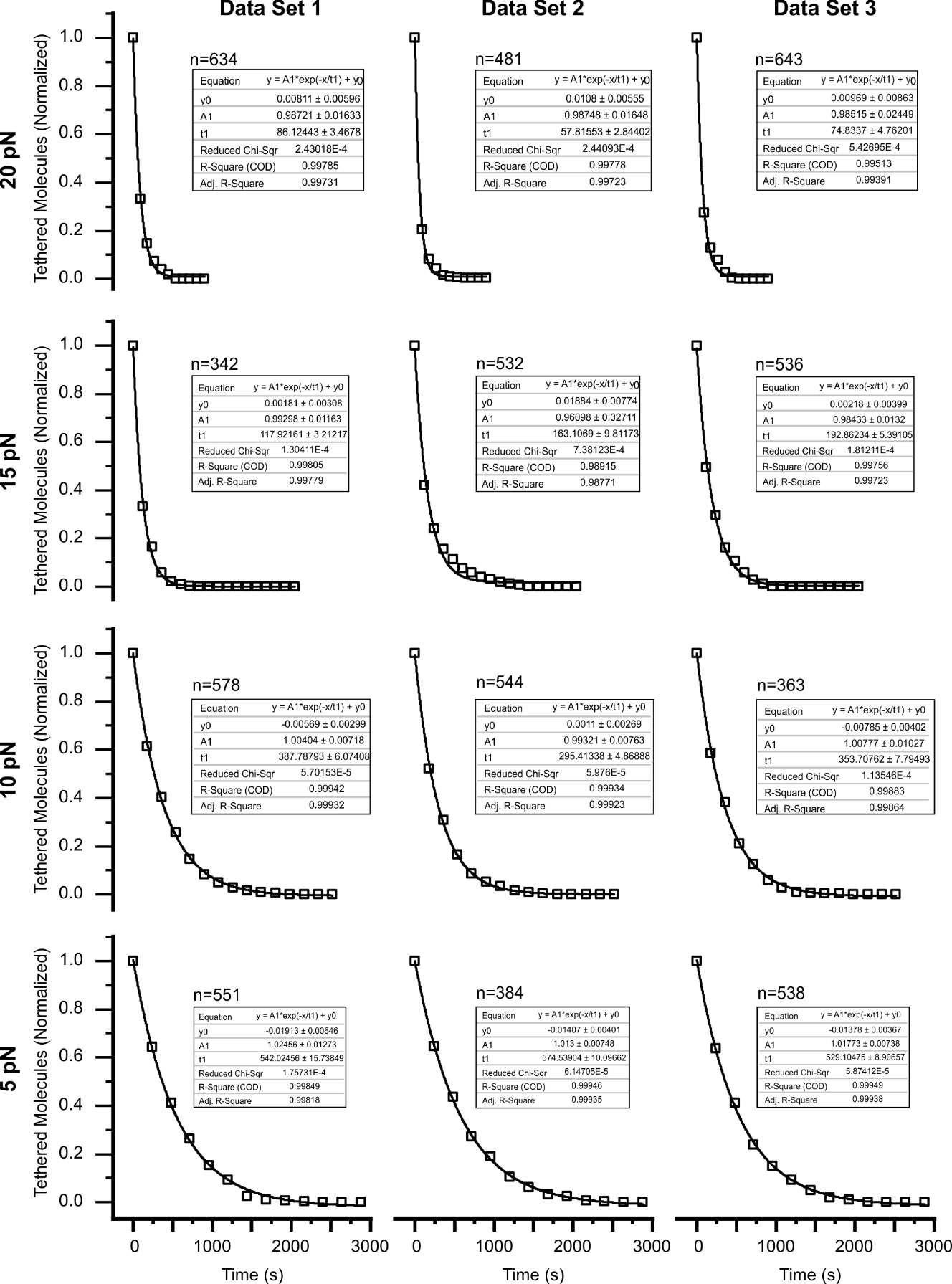

***Figure S3****:* ***Decay plot and single-exponential fitting of A|T construct at various forces.***

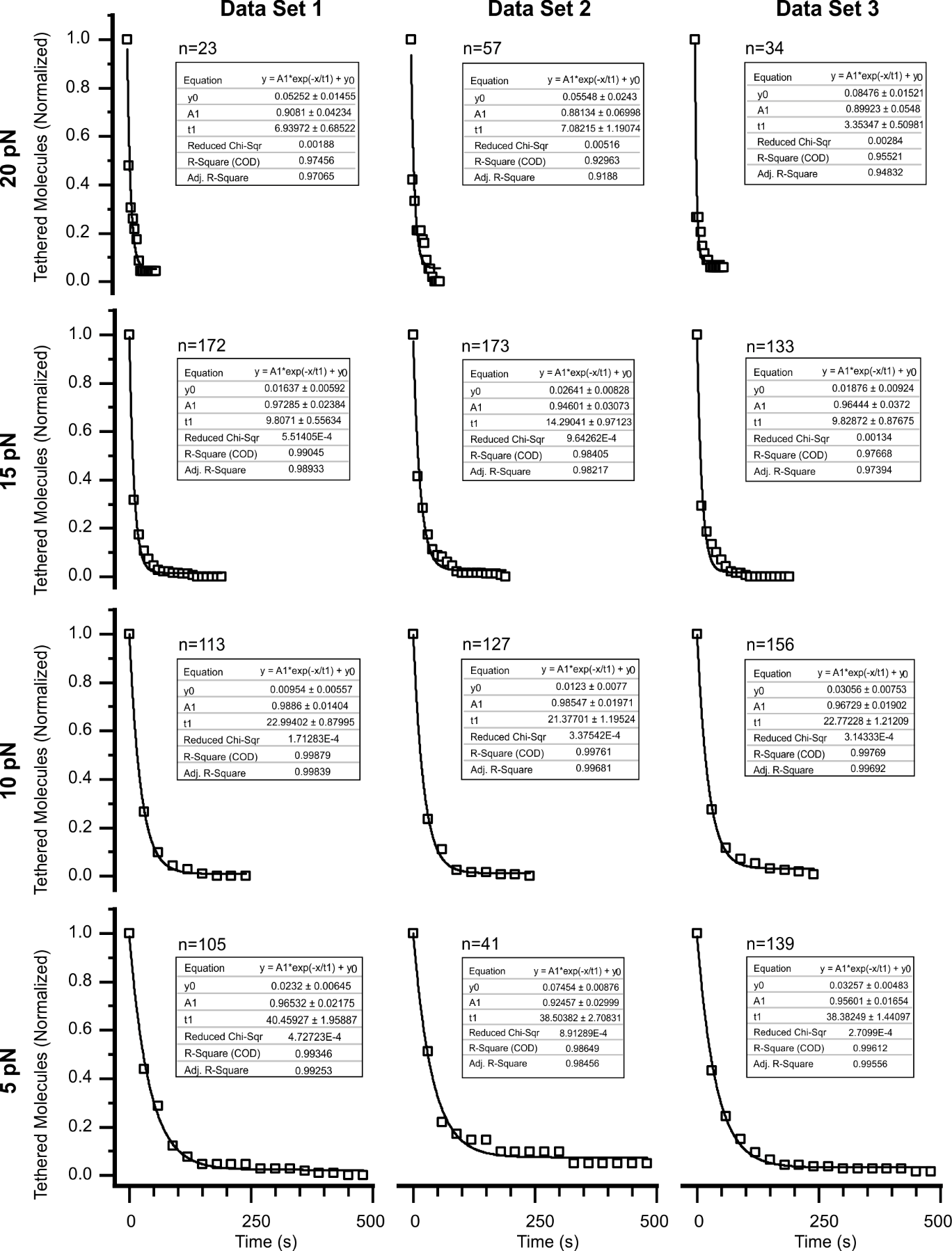

***Figure S4:*** ***Decay plot and single-exponential fitting of control construct at various forces.***

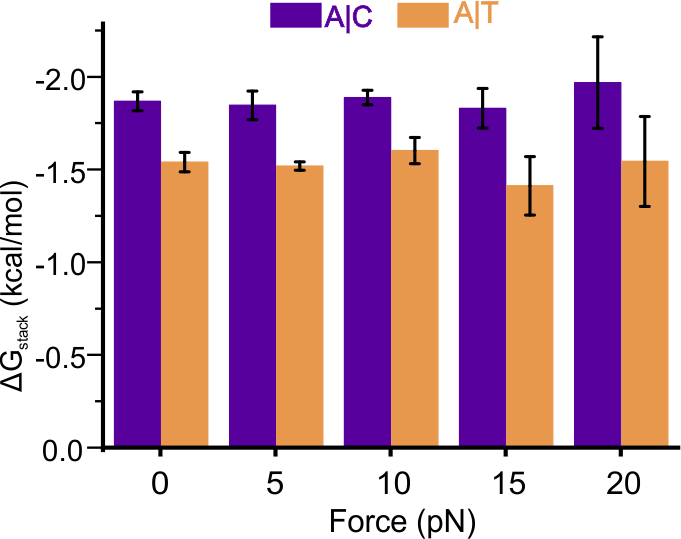

***Figure S5:*** ***ΔG_stack_ determined for A|C and A|T base-stack at various forces and at thermal equilibrium.***

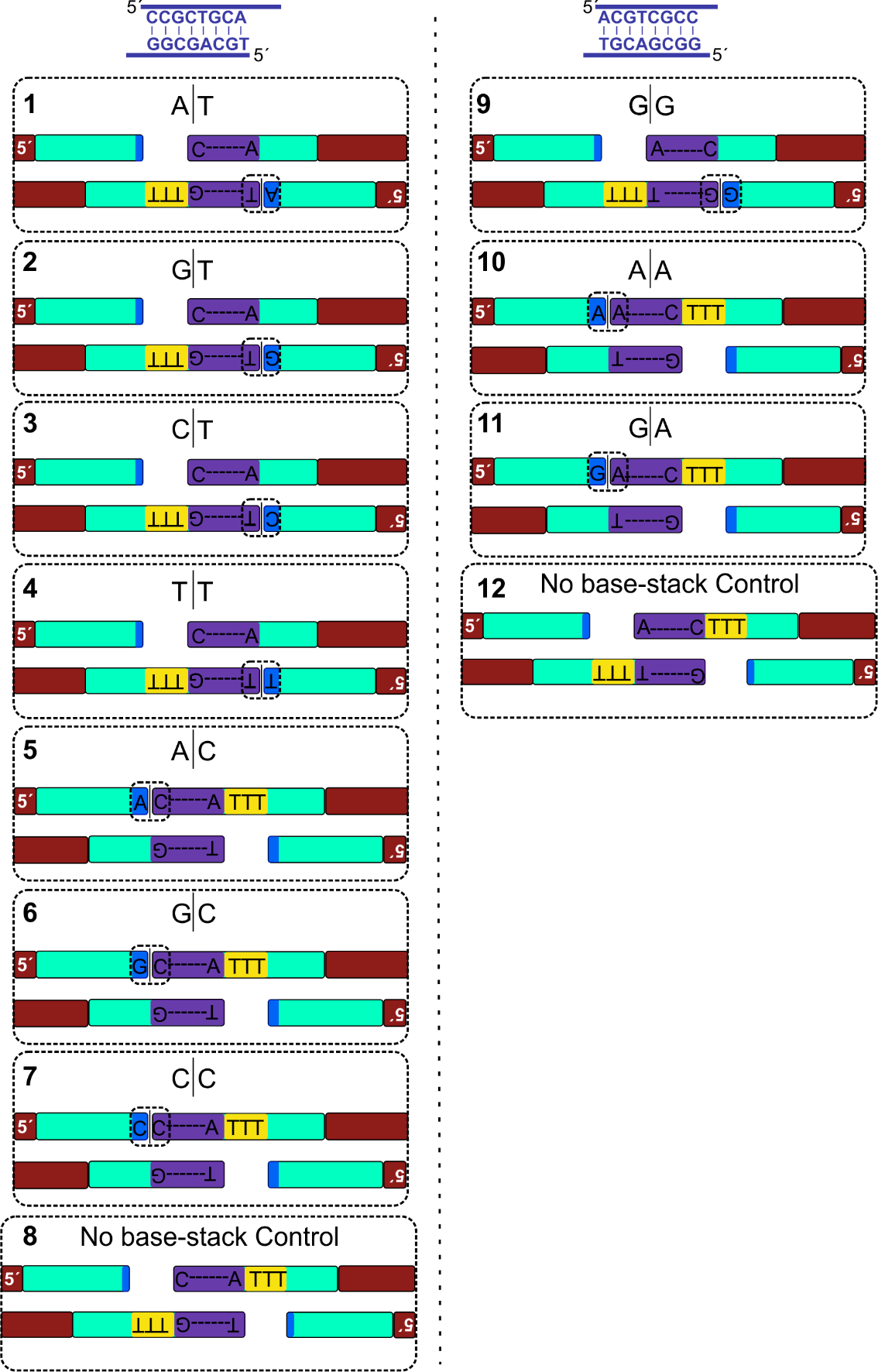

***Figure S6****:* ***Tethering combinations for all combinations.*** *The central duplex is conserved in all cases, but polarity is revered for the constructs in right to achieve all the combinations. The control constructs with both polarity is shown below on both left and right panel. The number on the top left of each box represents tether ID same as in Table S3.*

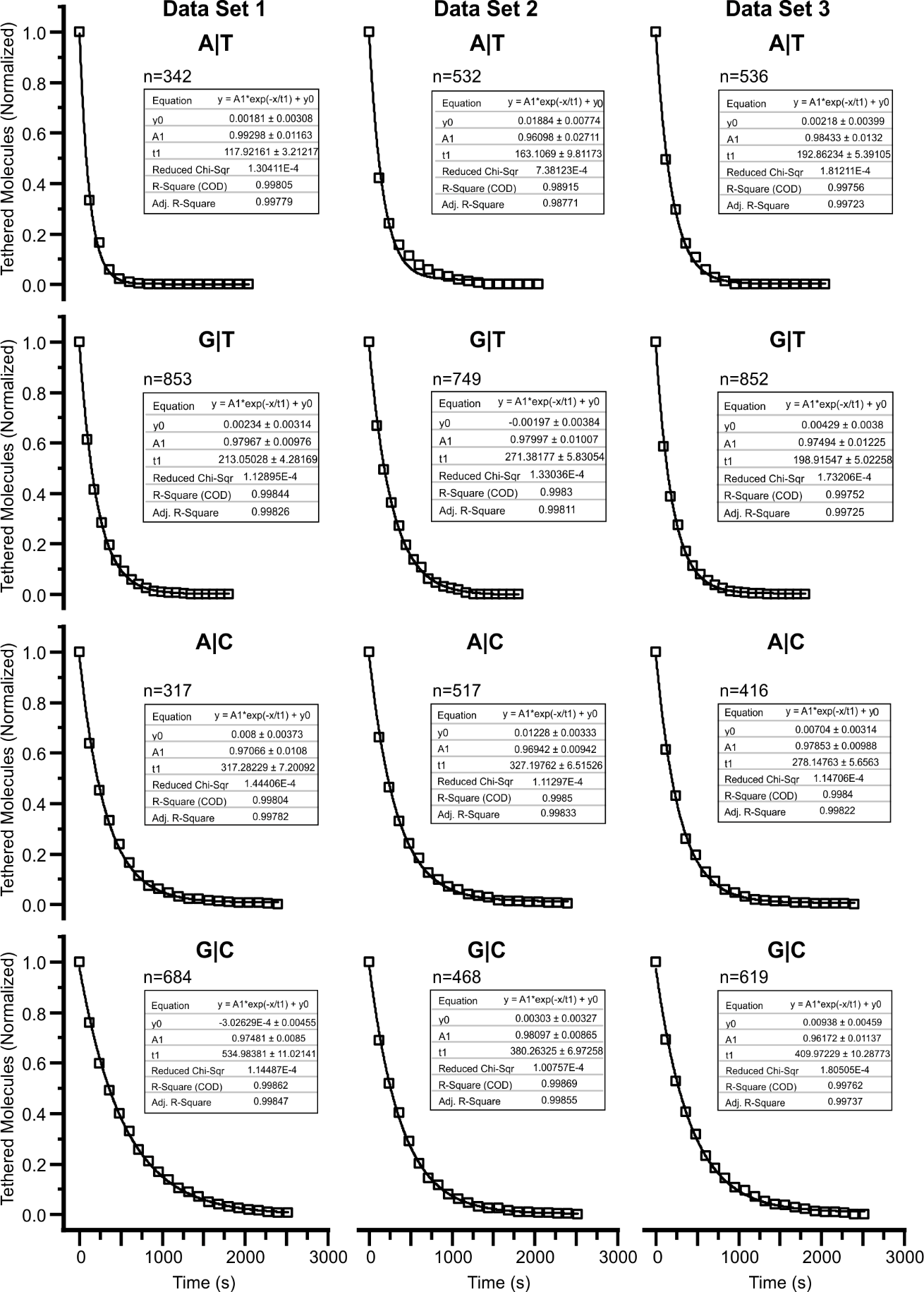

***Figure S7****:* ***Decay plot and single-exponential fitting of A|T, G|T, A|C and G|C combinations of base-stacks at 15 pN.***

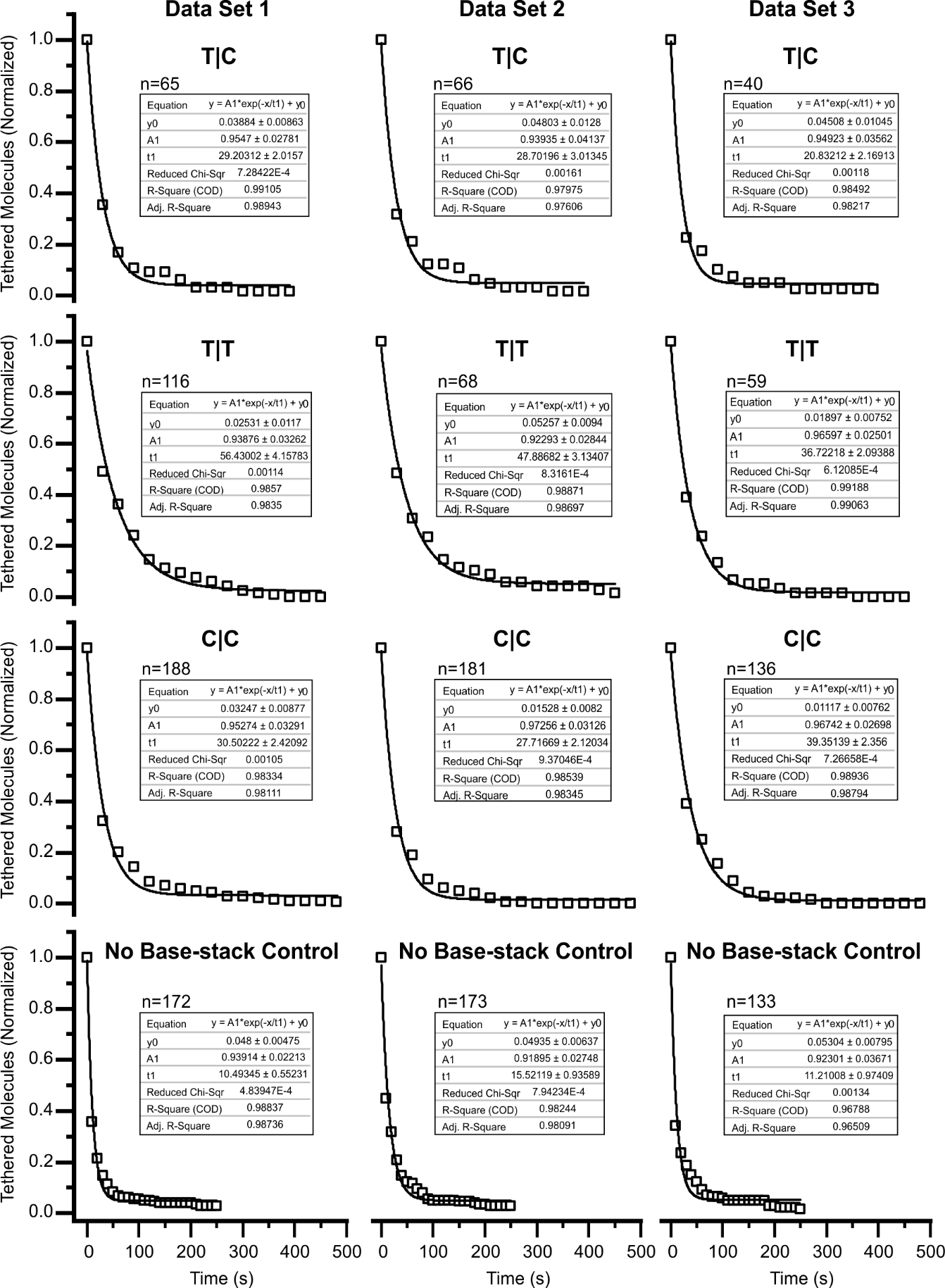

***Figure S8****:* ***Decay plot and single-exponential fitting of C|T, T|T, C|C combinations and control construct at 15 pN.***

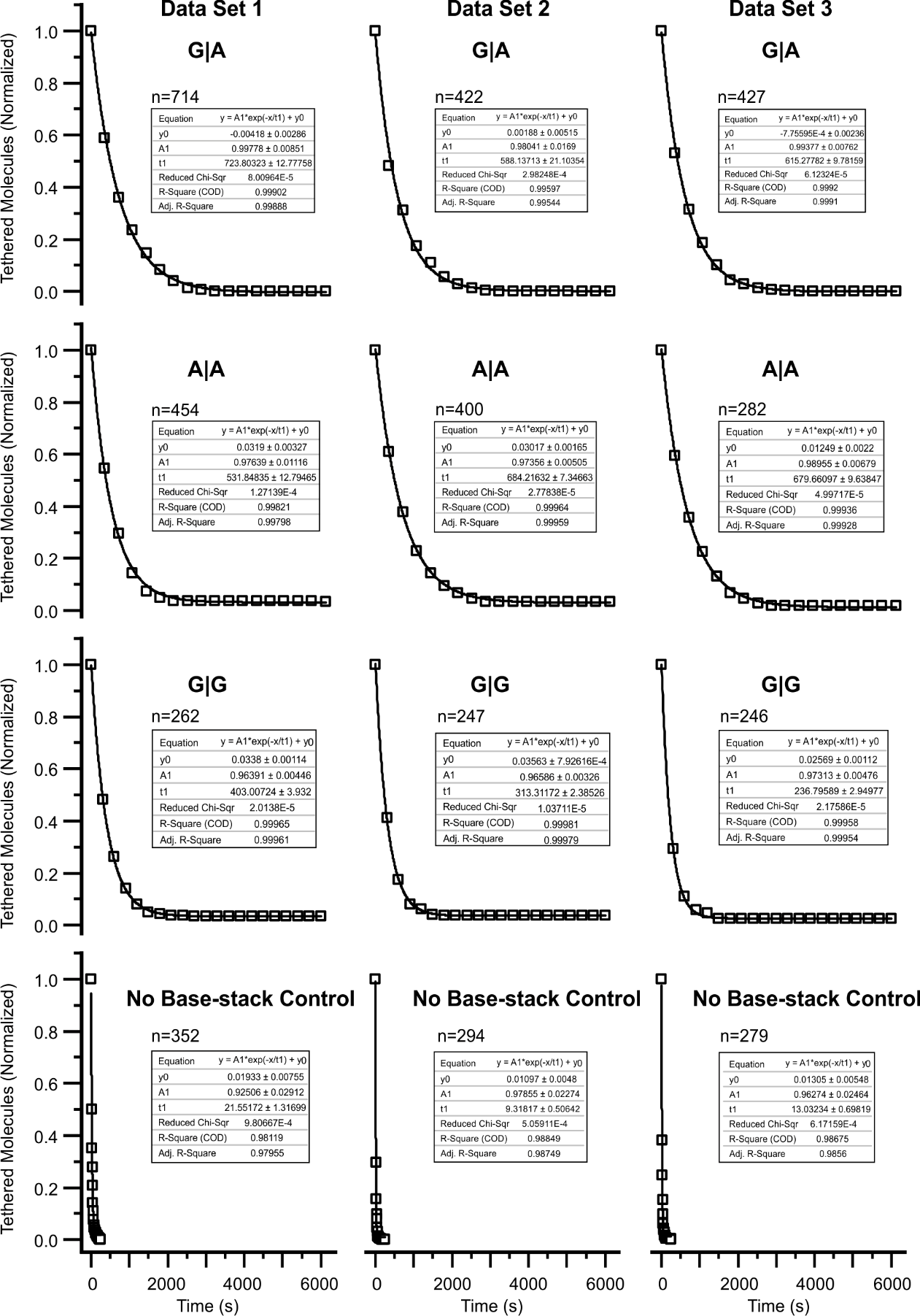

***Figure S9****:* ***Decay plot and single-exponential fitting of G|A, A|A, G|G combinations of base-stacks corresponding control constructs at 15 pN.***

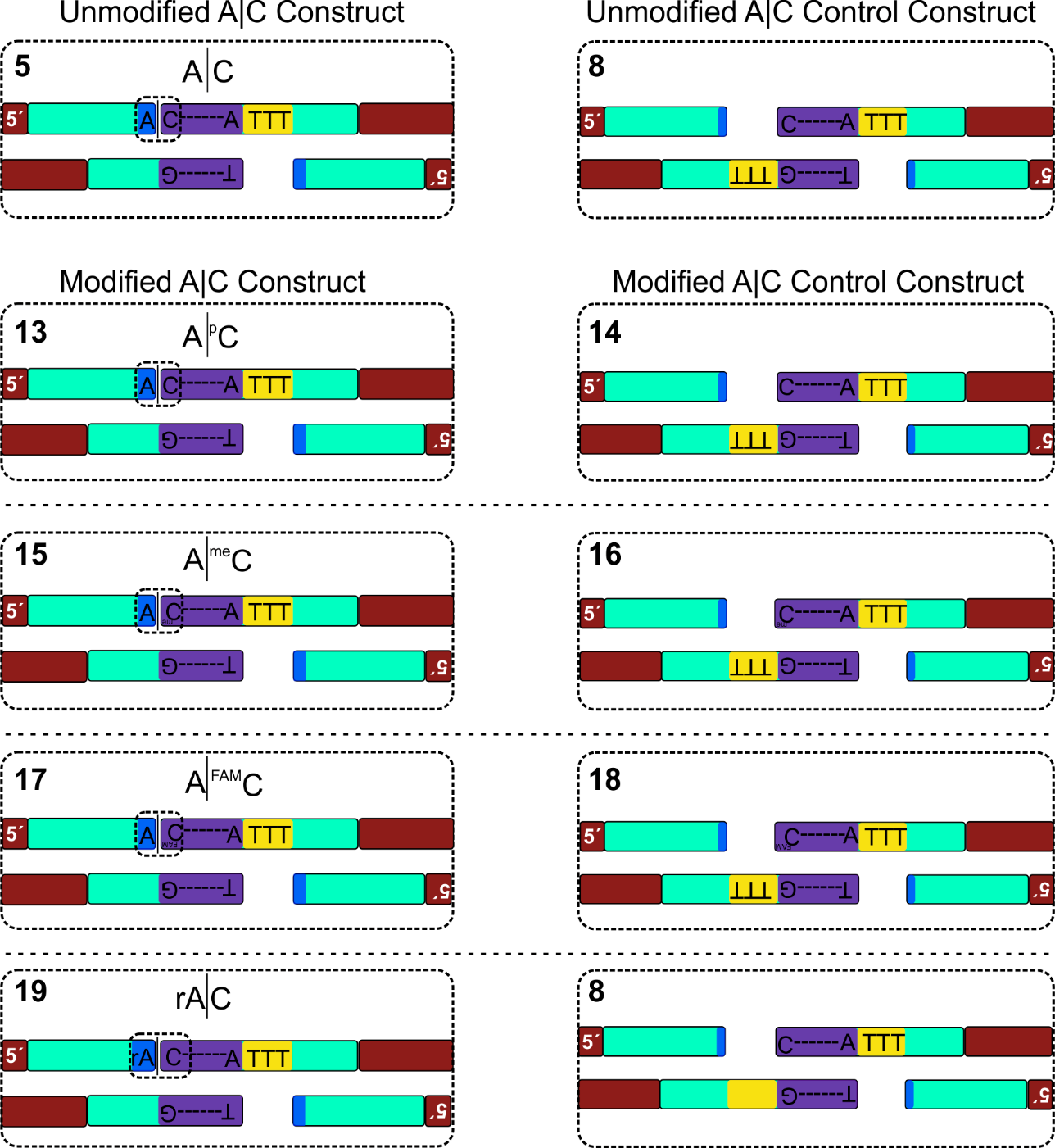

***Figure S10****:* ***Tethering combinations for modified base constructs.*** *The number on the top left of each box represents tether ID same as in Table S3.*

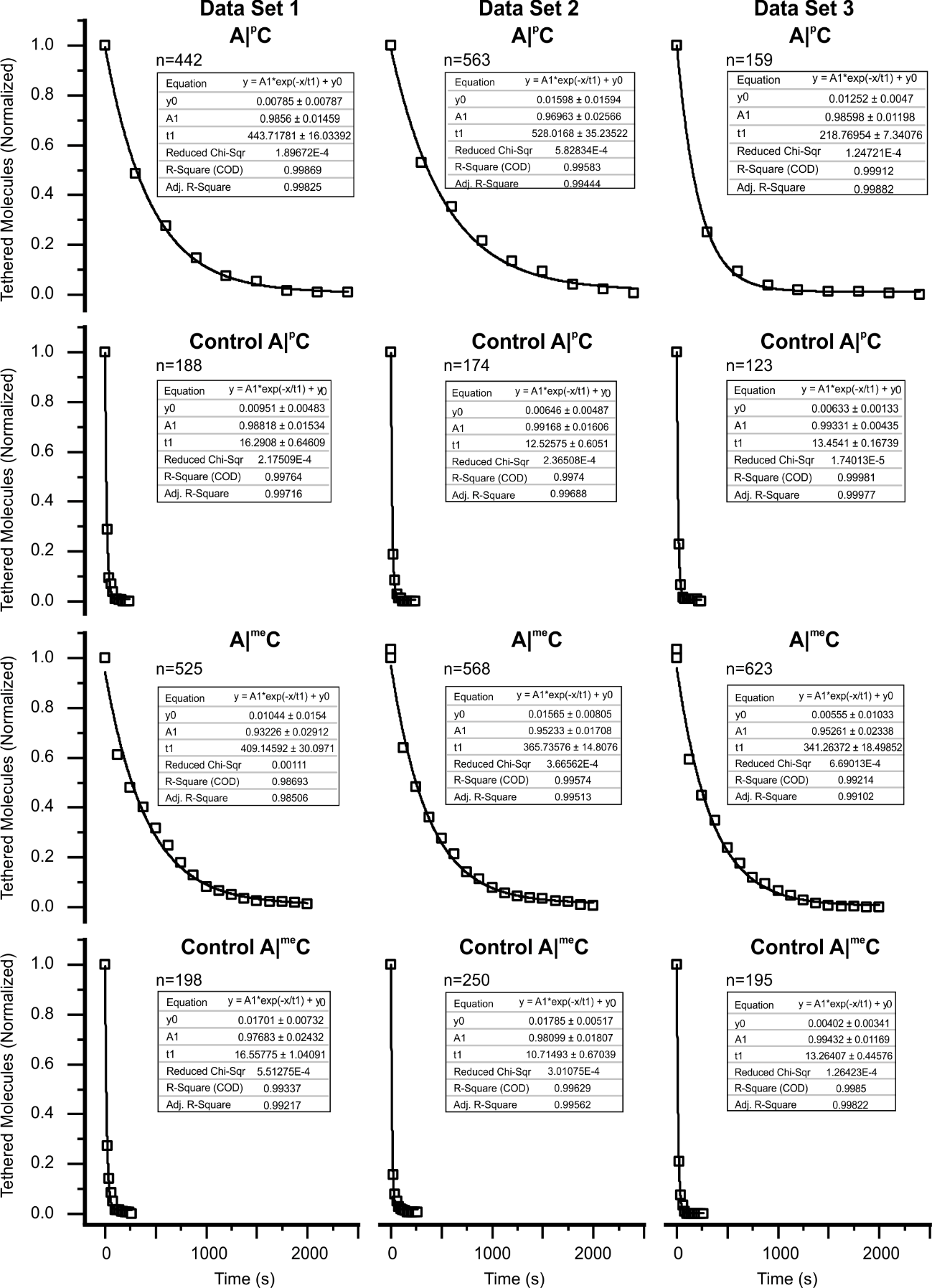

***Figure S11****:* ***Decay plot and single-exponential fitting of phosphorylated and methylated A|C base-stacks and their corresponding control constructs at 15 pN.***

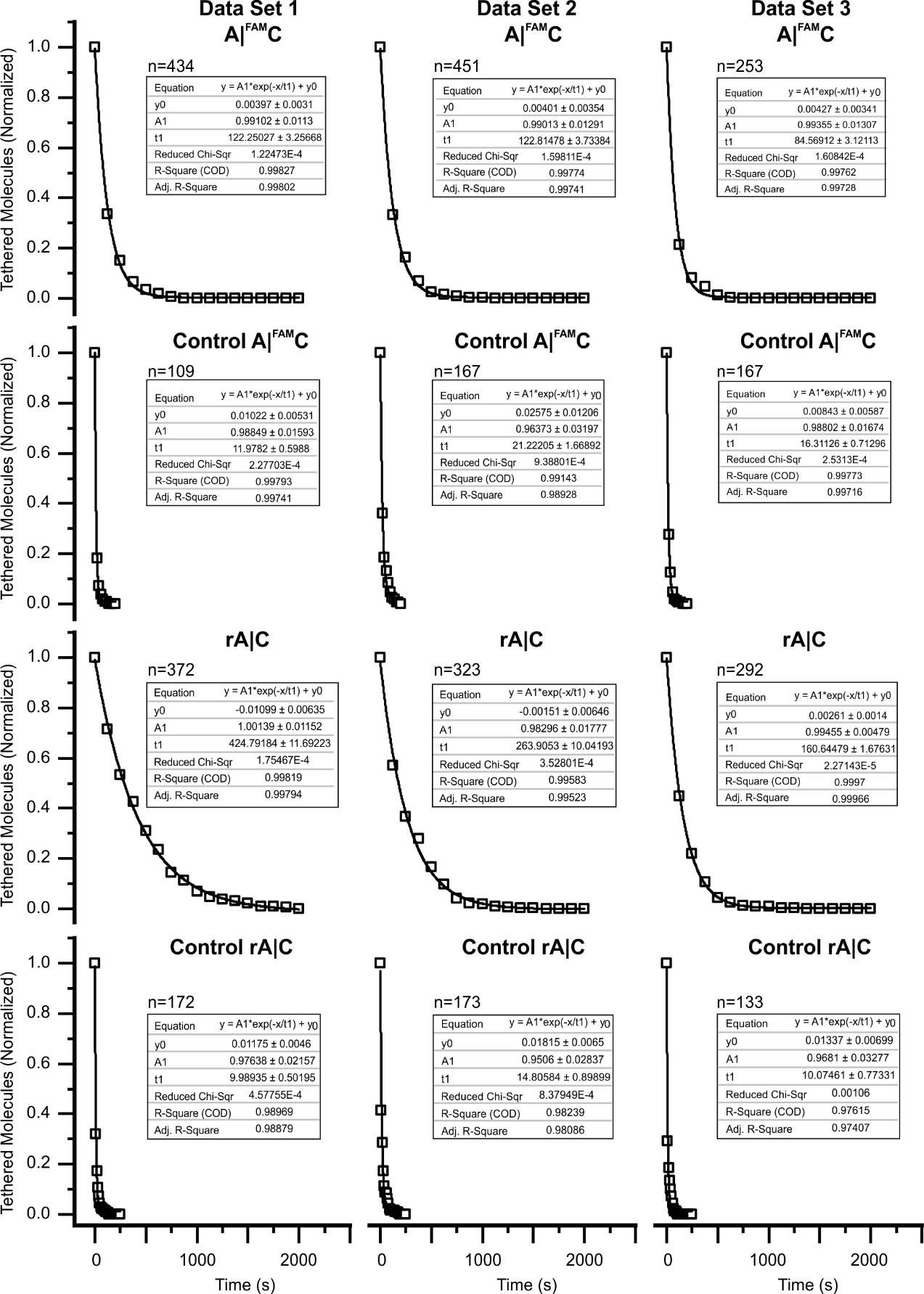

***Figure S12****:* ***Decay plot and single-exponential fitting of FAM and ribose A|C base-stacks and their corresponding control constructs at15 pN.***

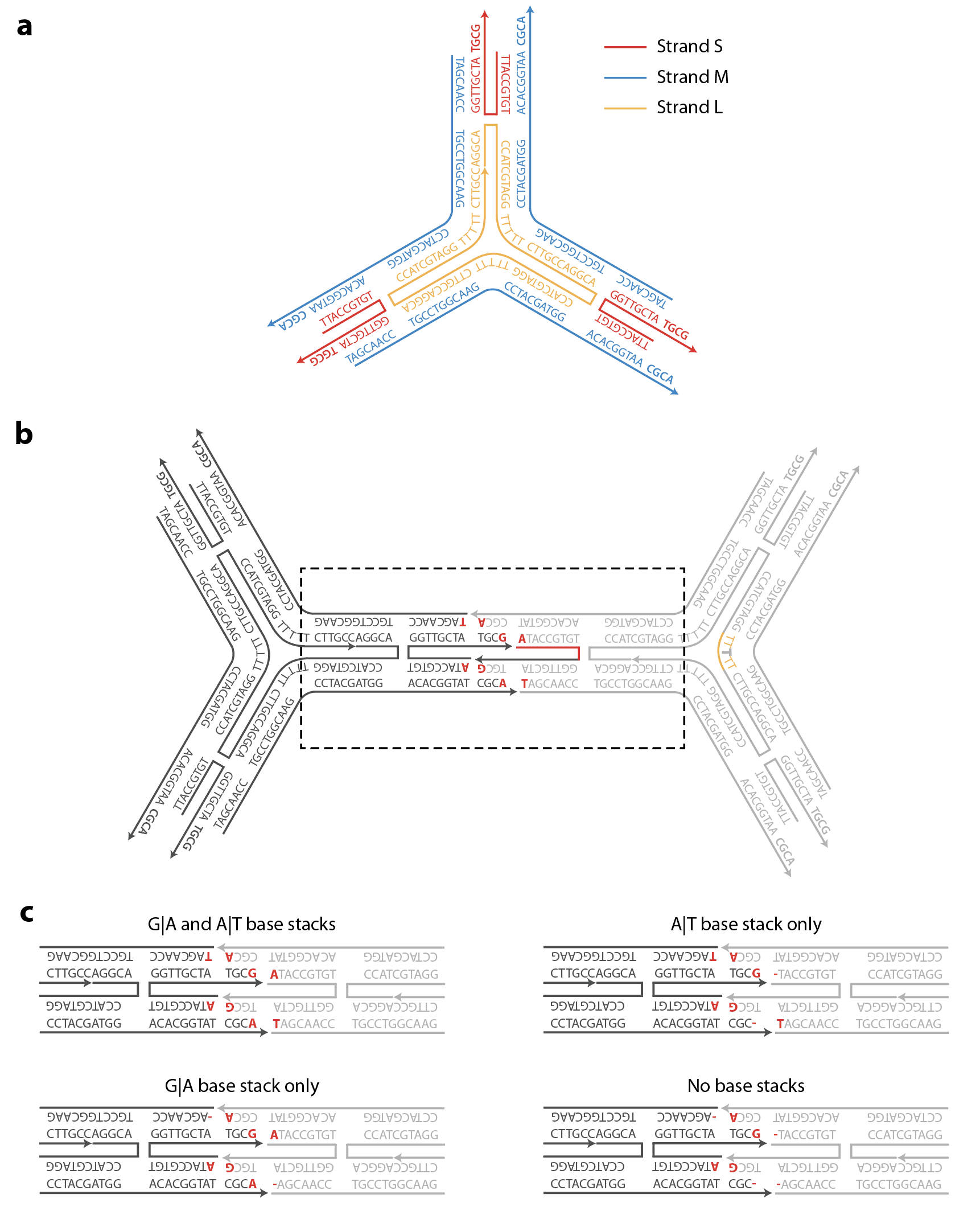

***Figure S13****:* ***Design of DNA tetrahedron.*** *(a) Design and sequences of the 3-point-star motif. (b) Illustration showing the interface of double cohesion with two sticky ends and corresponding base stacks. (c) Illustration showing the different base stack combinations tested in this study.*

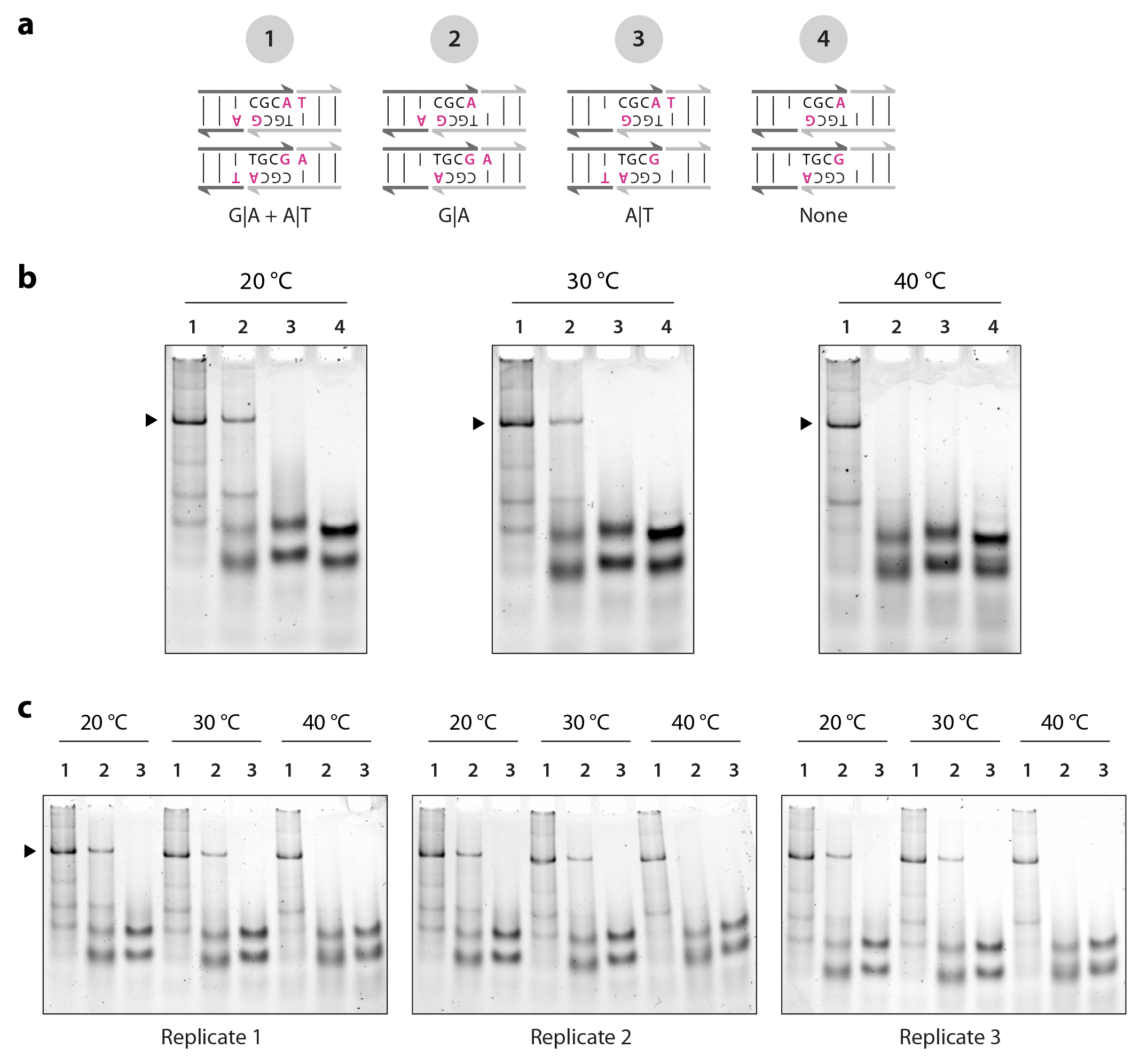

***Figure S14****:* ***Triplicate gel pics for DNA tetrahedron at various temperatures.***

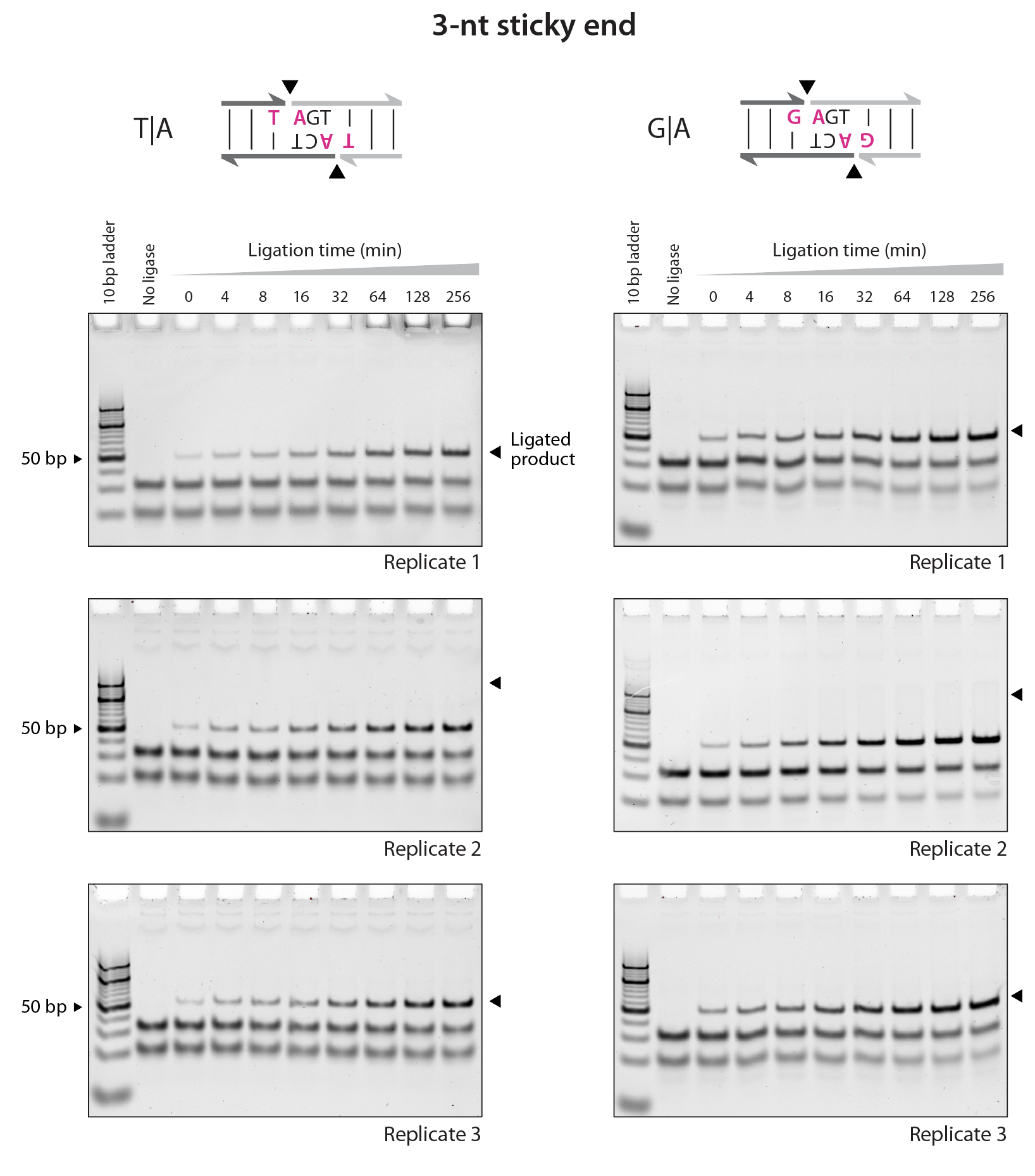

***Figure S15:*** ***Triplicate gel pics for ligation experiments with 3 nt overhang.***

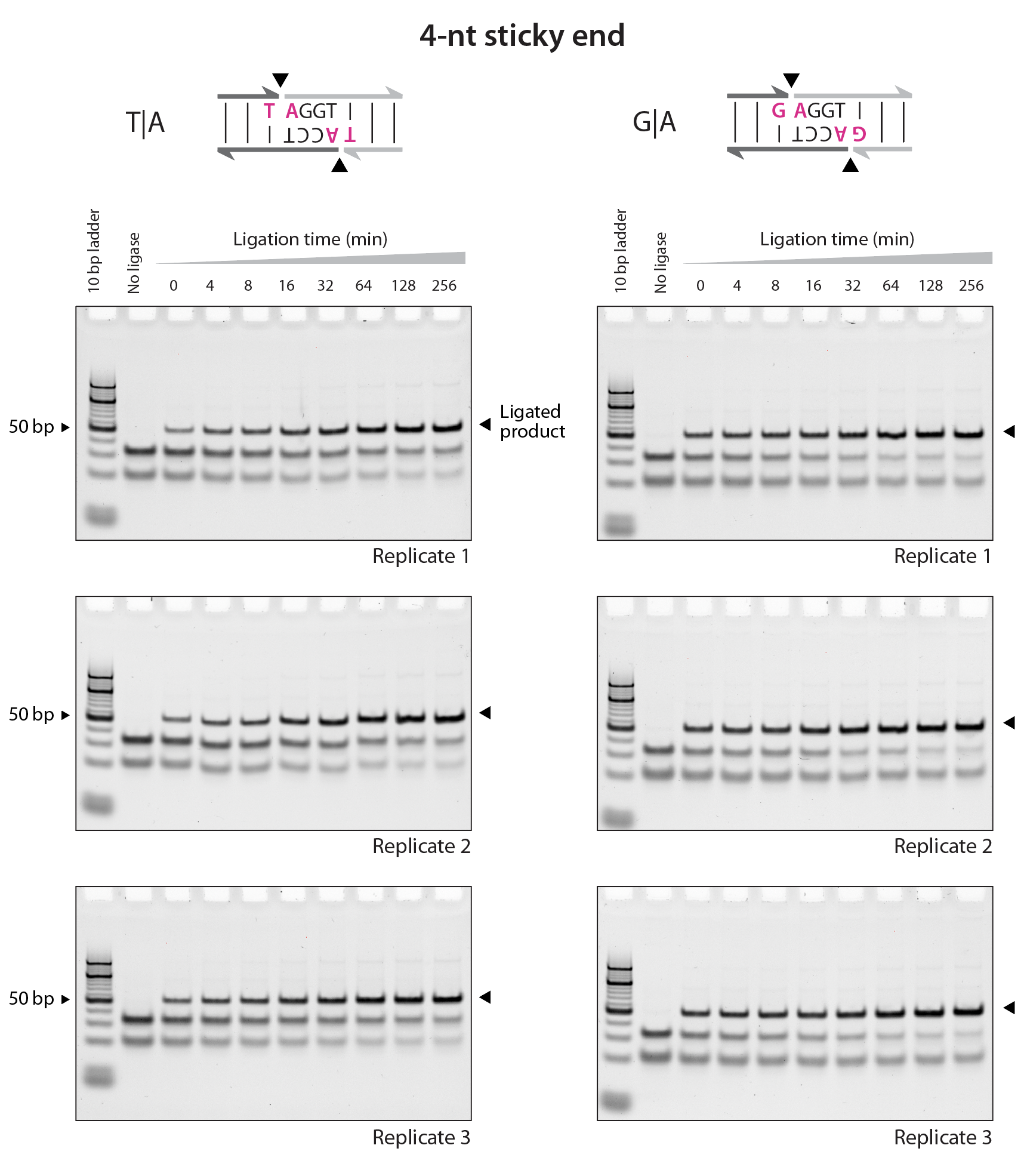

***Figure S16:*** ***Triplicate gel pics for ligation experiments with 4 nt overhang.***

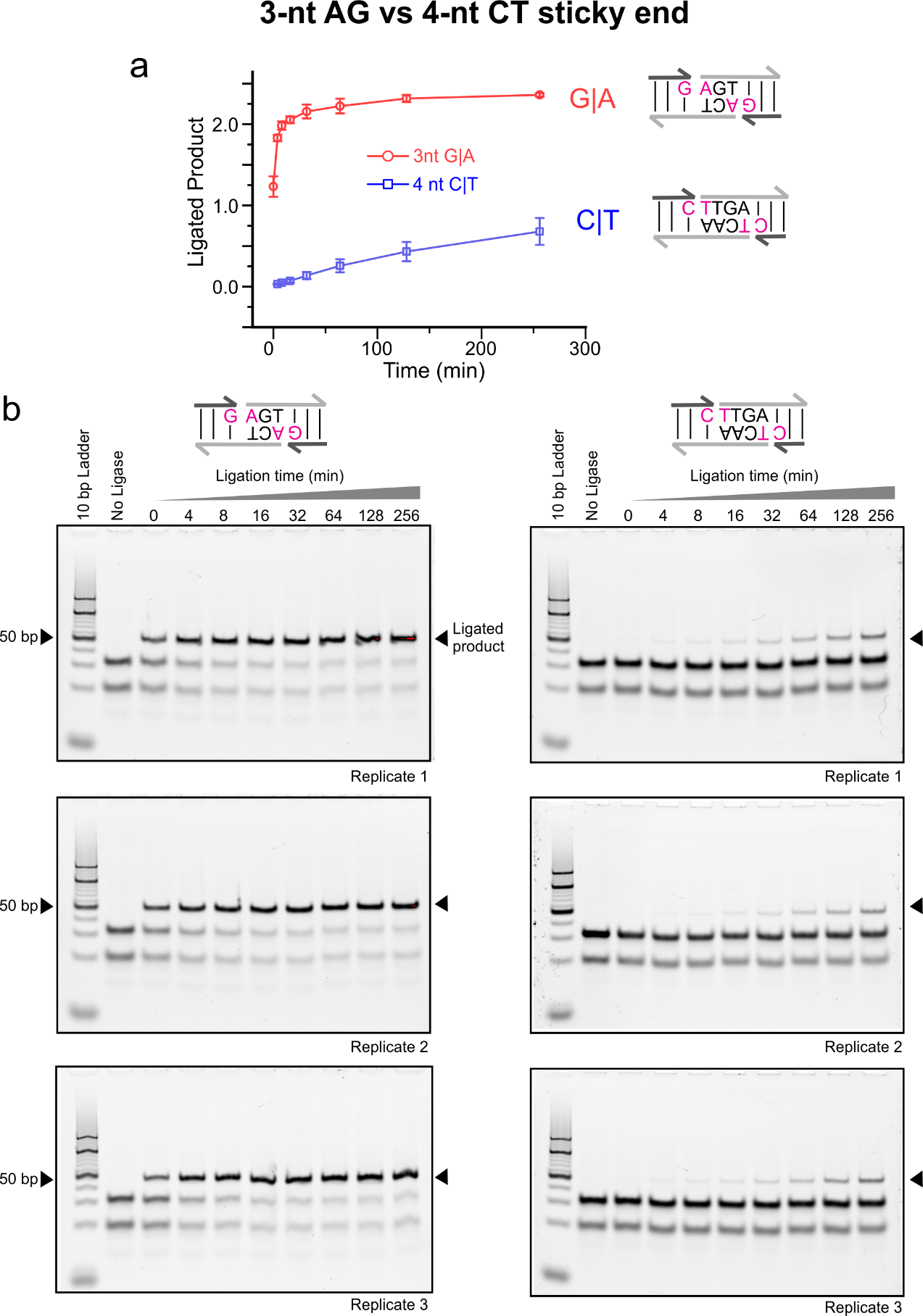

***Figure S17: Ligation experiments with 3 nt overhang with G|A Stack vs 4 nt overhang with C|T stack.*** *(a) Quantified ligation product of G|A and C|T stacked constructs over time. (b) Triplicate gel pictures.*
